# Ammonia inhibits antitumor activity of NK cells by decreasing mature perforin

**DOI:** 10.1101/2023.11.20.567708

**Authors:** Joanna Domagala, Tomasz M. Grzywa, Iwona Baranowska, Aleksandra Kusowska, Klaudyna Fidyt, Katsiaryna Marhelava, Zofia Pilch, Agnieszka Graczyk-Jarzynka, Lea K. Picard, Kamil Jastrzebski, Monika Granica, Magdalena Justyniarska, Doris Urlaub, Malgorzata Bobrowicz, Marta Miaczynska, Carsten Watzl, Magdalena Winiarska

**Affiliations:** Department of Immunology, Medical University of Warsaw, Warsaw, Poland; Laboratory of Immunology, Mossakowski Medical Research Institute, Polish Academy of Sciences, Warsaw, Poland; Doctoral School, Medical University of Warsaw, Warsaw, Poland; Department for Immunology, Leibniz Research Centre for Working Environment and Human Factors (IfADo) at TU, Dortmund, Germany; International Institute of Molecular and Cell Biology, Warsaw, Poland

**Author notes:** These authors contributed equally: Joanna Domagala, Tomasz M. Grzywa. The current address: The Raymond G. Perelman Center for Cellular and Molecular Therapeutics, Department of Pathology and Laboratory Medicine, Children’s Hospital of Philadelphia, Philadelphia, USA. Correspondence to: Magdalena Winiarska, PhD.

## Abstract

Immunotherapy revolutionized cancer treatment in the last decade. Natural killer (NK) cells are one of the key host immunity components against malignant cells. Thus, they are currently extensively investigated in the field of immunotherapy of cancer. Different approaches have been developed to improve the antitumor activity of NK cells. Nonetheless, tumor microenvironment remains an obstacle to effective NK cell-based therapies. Here, we demonstrated that a cancer-conditioned medium suppresses the anti-tumor activity of NK cells. Further, we found that ammonia, a by-product of cancer cell metabolism, accumulates in the cancer-conditioned medium and tumor microenvironment. We identified that ammonia impairs the cytotoxicity of NK cells as well as the effectiveness of antibody-based and chimeric antigen receptor (CAR)-NK-based therapies *in vitro*. Inhibited activity of NK cells was caused by decreased levels of perforin. This effect was dependent on the lysosomotropic features of ammonia and its ability to increase pH in acidic compartments. In consequence, upon contact with ammonia the mature form of perforin was decreased in NK cells leading to their dysfunction. Our findings demonstrate that in addition to its previously described role of promoting tumor growth as a nitrogen source for tumor biomass ammonia could promote tumor escape as an NK cells immune checkpoint.

**Graphical abstract:** 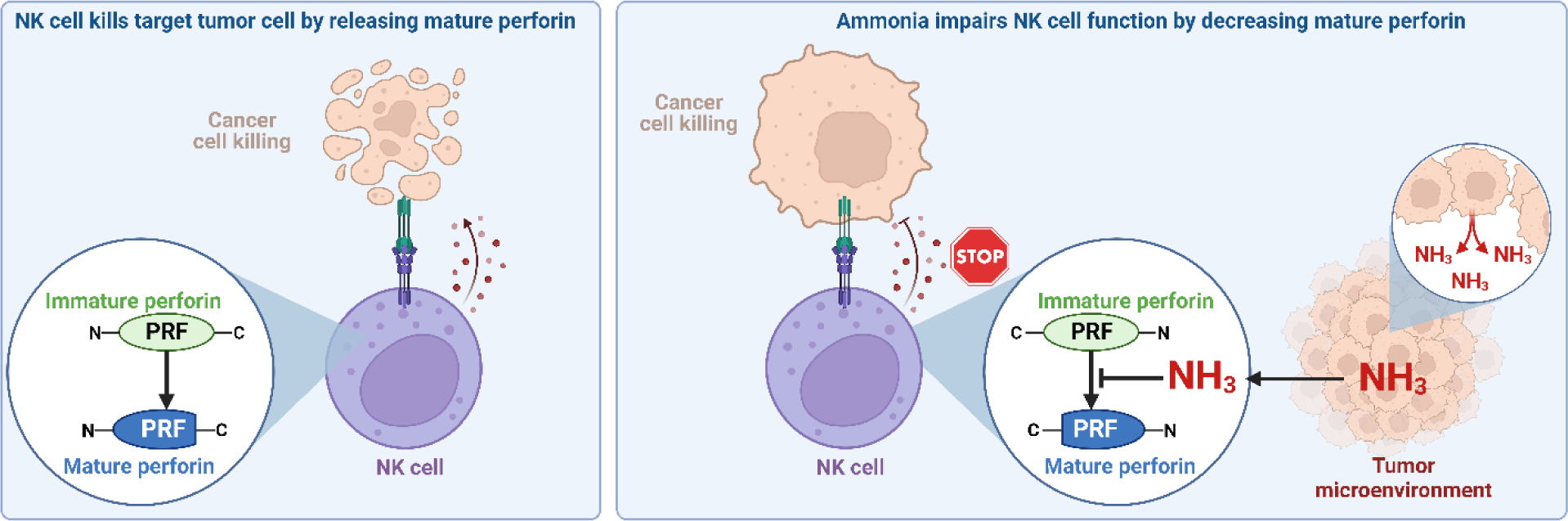

**Highlights:**

- Cancer-conditioned medium suppresses the antitumor activity of NK cells
- Ammonia accumulates in conditioned medium and in the tumor microenvironment
- Impaired cytotoxicity of NK cells is caused by ammonia that decreases perforin levels
- Ammonia causes NK cell dysfunction

## INTRODUCTION

Natural killer (NK) cells constitute one crucial component of the antitumor immune response ^1^. Due to their potent cytotoxic activity against malignant cells as well as relatively low sensitivity to inhibitory signals in cancer, they are extensively studied in the field of immunotherapy. Beyond their natural cytotoxic potential, NK cells can kill tumors in a mechanism of antibody-dependent cell-mediated cytotoxicity (ADCC) by engagement with monoclonal antibodies, which include clinically available antibodies targeting CD20 (rituximab, RTX), CD38 (daratumumab), and HER2 (trastuzumab)^2^. Additionally, NK cells can be used in adoptive cell therapies as chimeric antigen receptor-modified (CAR)-NK cells and are a promising alternative to CAR-T cells ^3^. Nonetheless, multiple factors secreted by cancer cells and other types of cells in the tumor microenvironment suppress their activity, including not only protein factors like cytokines but also different metabolites ^4,5^.

Dysregulation of cellular metabolism is one of the hallmarks of cancer ^6^. For instance, cancer cells rely on glycolysis and glutaminolysis to a much greater extent than healthy cells ^7^. This dysregulation results in the specific profile of metabolites in the tumor microenvironment (TME) ^8^. While in normal tissues toxic end products of metabolism are excreted, impaired tumor tissue architecture and abnormal vascularization lead to their accumulation in TME. That includes lactate and ammonia which promote tumor growth by serving as energy and nitrogen source, respectively ^9,10^.

Recently, ammonia concentrations have been demonstrated to be substantially increased TME where immune cells interact with the cancer cells. Previous studies also revealed that ammonia may regulate immune response by suppressing the activity of macrophages, dendritic cells, and neutrophils ^11–15^. Furthermore, ammonia has been identified as a driver of T cell exhaustion and a regulator of the development of memory T cells ^16–18^. However, whether ammonia impairs the cytotoxic functions of immune cells, including NK cells, remains unknown.

In this study, we demonstrated that a cancer-conditioned medium suppresses the antitumor activity of NK cells. We identified that ammonia accumulates in cancer-conditioned medium and tumor interstitial fluid (TIF) in mice and inhibits the cytotoxic activity of NK cells by decreasing the amount of mature perforin in secretory lysosomes. Collectively, our findings suggest that ammonia plays a role as a metabolic checkpoint for NK cells.

## RESULTS

### Cancer-conditioned medium suppresses NK cells

To determine the impact of cancer-secreted factors on NK cell activity, we cultured different cell lines for 48 hours to generate a cancer-conditioned medium. Then, natural (spontaneous) cytotoxicity of NK cells was determined by their co-culture with K562 cells in the conditioned or control medium. We found that medium conditioned by various tumor cells such as human lymphoma cell lines (Raji, Daudi, and Ramos cells, Fig. 1a), human multiple myeloma cell lines (MM.1s, H929, Fig. 1b), breast cancer cell lines (human HCC1806, murine 4T1, E0771, EMT6) (Fig. 1c) significantly inhibited natural cytotoxicity of NK cells against K562 cells. In contrast, medium conditioned by non-malignant fibroblast cell line (L929) and activated human T cells had no effect on the cytotoxicity of NK cells (Fig. 1d).

**Fig. 1.**
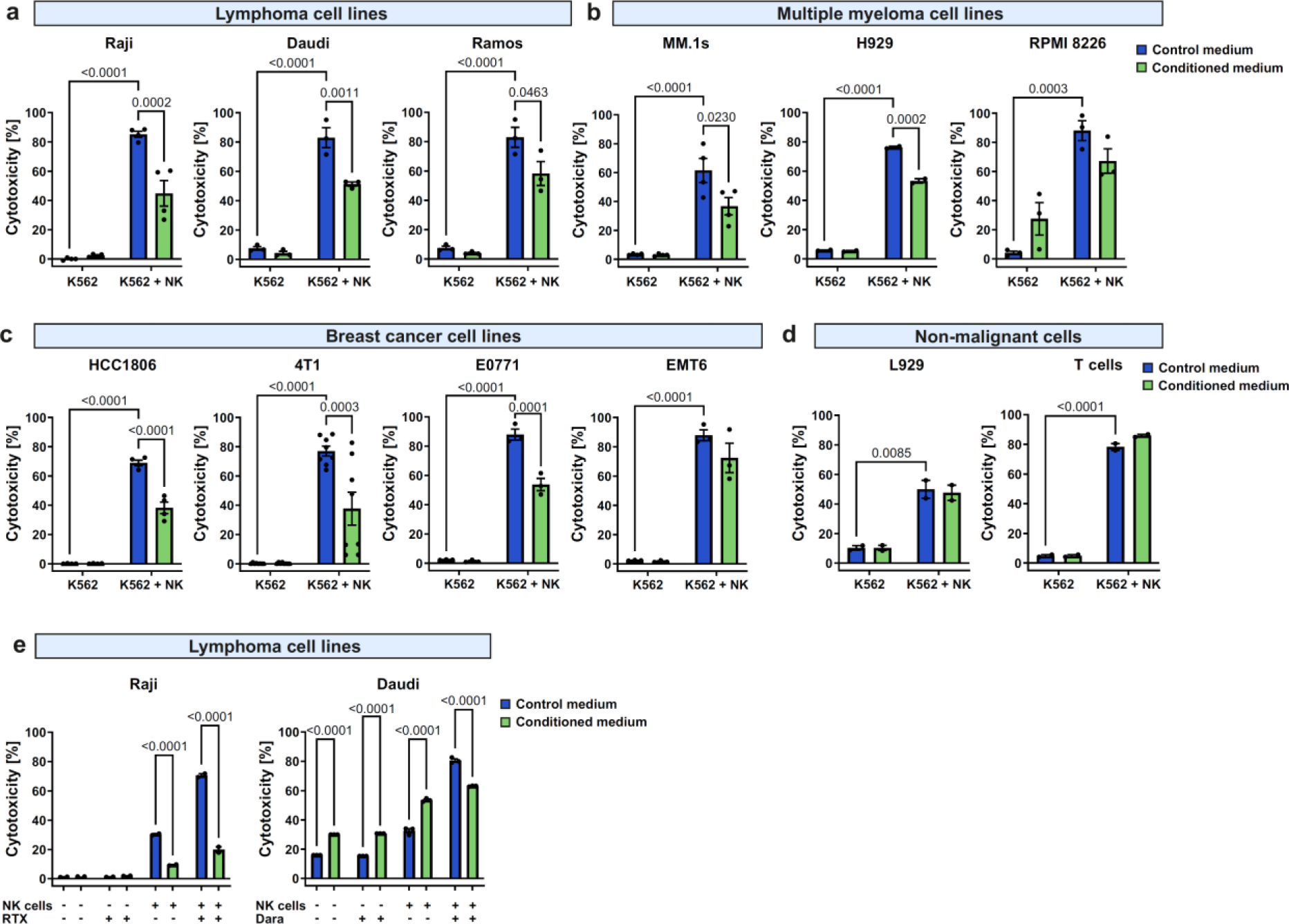
Cancer cell-conditioned medium inhibits natural cytotoxicity and ADCC of NK cells. **a**, Natural cytotoxicity of NK cells against K562 cells in the presence of control medium and lymphoma cells-conditioned medium (Raji, n=4; Daudi, n=3; Ramos, n=3). **b**, Natural cytotoxicity of NK cells against K562 cells in the presence of control medium and multiple myeloma cells-conditioned medium (MM.1s, n=4; H929, n=2; RPMI 8226, n=3). **c**, Natural cytotoxicity of NK cells against K562 cells in the presence of control medium and breast cancer cells-conditioned medium (HCC1806, n=4; 4T1, n=8; E0771, n=3; EMT6, n=3;). **d**, Natural cytotoxicity of NK cells against K562 cells in the presence of control medium and non-malignant cells-conditioned medium (L929, n=2; T-cells, n=2). For **a-d**, K562 cells were stained with CFSE and incubated with NK cells in a medium conditioned by indicated cells. Cytotoxicity was assessed after 4 hours using flow cytometry as percentage of propidium iodide-positive CFSE-positive (K562) cells. **e**, ADCC of NK cells against Raji cells with anti-CD20 antibody (100 µg/ml of rituximab, RTX) and Daudi cells with anti-CD38 antibody (1 µg/ml of daratumumab, Dara) in the presence of control medium or lymphoma cells-conditioned medium (n=2). Cytotoxicity was assessed after 4 hours using flow cytometry as percentage of propidium iodide-positive CFSE-positive cells. *P* values were calculated using two-way ANOVA with Tukey’s post hoc test. Data show individual values and means ± SEM. *n* values are the numbers of biological replicates in *in vitro* experiments. The source data underlying **a-e** are provided as a Supplementary Data file.

NK cells may be engaged by monoclonal antibodies to mediate antitumor response ^19^. Thus, we subsequently determined whether the cancer-conditioned medium affected the antibody-dependent cell-mediated cytotoxicity (ADCC) of NK cells. For this purpose, we employed lymphoma and multiple myeloma cell line models, expressing CD20 and CD38, respectively, as targets for clinically used antibodies. We found that tumor cells-conditioned medium suppressed RTX- and daratumumab-dependent cell-mediated cytotoxicity of NK cells against CD20^+^ Raji cells and CD38^+^ Daudi cells, respectively (Fig. 1e). Further, to recapitulate natural architecture of solid tumor more reliably, including heterogeneous distribution of oxygen and nutrients differently affecting cell metabolism, we utilized a three-dimensional (3D) cell culture of both murine (4T1) and human (MDA-MB 231, HCC1806) breast cancer cells (Supplementary Fig. 1a) and found out that conditioned medium potently inhibited cytotoxicity of NK cells (Supplementary Fig. 1b).

To characterize factors responsible for the suppression of NK cells, we fractionated a representative cancer-conditioned medium from Raji cells using ultrafiltration into fractions containing factors smaller than 3 kDa (predominantly low-molecular weight metabolites) and larger molecules (predominantly proteins) (Supplementary Fig. 2a). It revealed that only low molecular-weight fraction of Raji cells-conditioned medium impaired cytotoxicity of NK cells against target cells (Supplementary Fig. 2b).

### Ammonia accumulates in cancer cell-conditioned medium and TME

Based on the literature data that demonstrated the accumulation of ammonia in TME and its inhibitory effects on different populations of immune cells ^10,14,16,20^, we hypothesized that it may contribute to the inhibitory effects of low molecular-weight fraction of cancer-conditioned medium on NK cells activity. Indeed, accumulation of ammonia was detected in media conditioned by all checked tumor cell lines including lymphoma (Raji, Ramos, Daudi, Fig. 2a), multiple myeloma (MM.1s, RPMI 8226, H929, Fig. 2b), and breast cancer cell lines (MCF-7, EMT6, E0771 Fig. 2c).

**Fig. 2.**
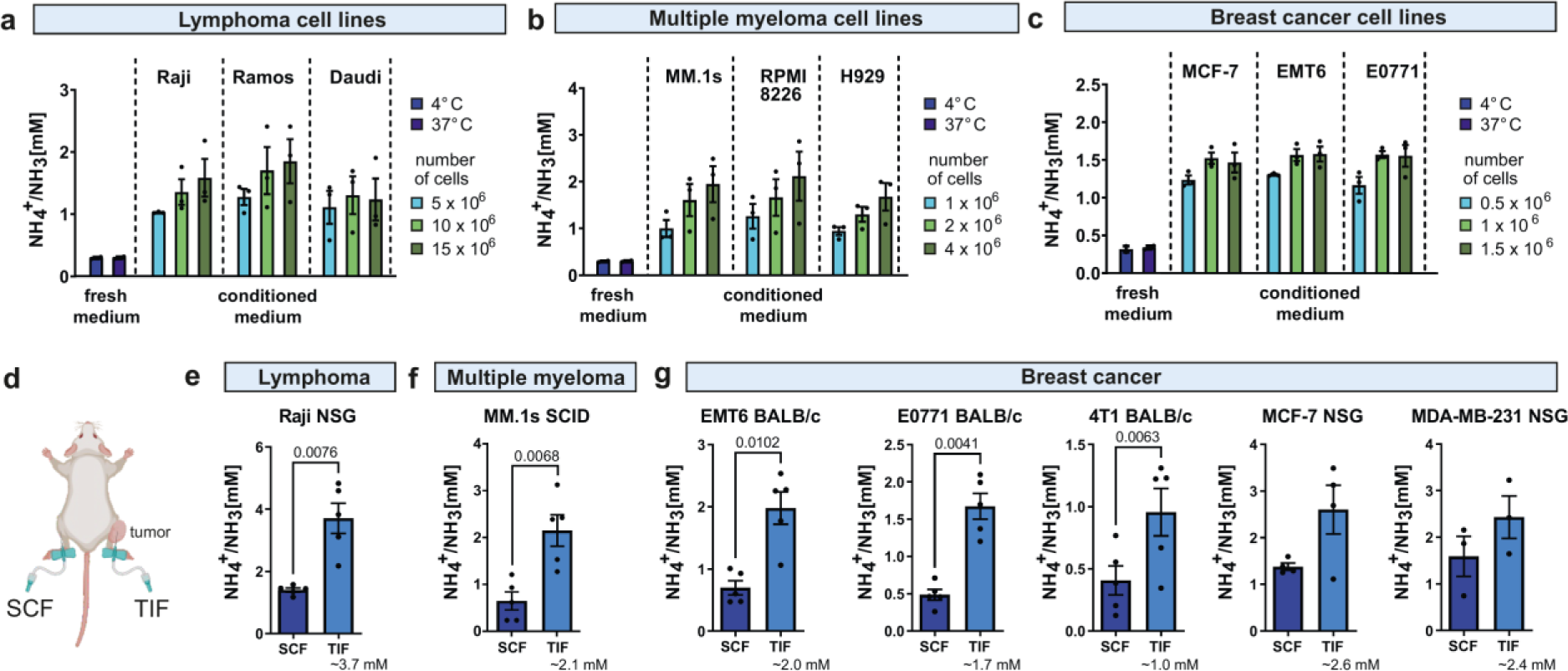
Ammonia concentration is increased in cancer cell-conditioned medium and tumor interstitial fluid (TIF) **a**, The concentration of ammonia in culture media after the culture of lymphoma cell lines for 48 hours (conditioned medium) in different densities (5.0 × 10^6^, 10.0 × 10^6^, 15.0 × 10^6^/ml) was measured using a Dimension Ammonia assay (Siemens) (n=3). **b**, The concentration of ammonia in culture media after culture of multiple myeloma cell lines for 48 hours (conditioned medium) in different densities (1 × 10^6^, 2 × 10^6^, 4 × 10^6^/ml) measured using a Dimension Ammonia assay (Siemens) (n=3). **c**, The concentration of ammonia in culture media after culture of breast cancer cell lines for 48 hours (conditioned medium) in different densities (0.5 × 10^6^, 1 × 10^6^, 1.5 × 10^6^/ml) measured using a Dimension Ammonia assay (Siemens) (n=3). In **a-c** empty medium incubated for 48 hours (without the cells) at 4°C or 37°C is presented as control. **d**, Schematic presentation of the tumor interstitial fluid (TIF) and subcutaneous fluid (SCF) isolation from mice. TIF was collected from tumors not exceeding 1500 mm^3^. SCF was isolated at the same time from the contralateral tight as a control tissue fluid. **e**, The concentration of ammonia in TIF and SCF isolated from Raji tumor-bearing NSG (n=5). **f,** The concentration of ammonia in TIF and SCF isolated MM.1s tumor-bearing SCID mice (n=5). **g,** The concentration of ammonia in TIF and SCF isolated from breast cancer-bearing mice (EMT6 in BALB/c mice, n=5; E0771 in BALB/c mice n=5; 4T1 in BALB/c mice, n=5; MCF-7 in NSG mice, n=4; MDA-MB-231 in NSG mice, n=3). *P* values were calculated using a paired t-test. Data show individual values and means ± SEM. *n* values are the numbers of biological replicates in *in vitro* experiments or number of mice used to obtain the data. The source data underlying **a-c, e-g** are provided as a Supplementary Data file.

To determine whether ammonia accumulation also occurs *in vivo,* we isolated tumor interstitial fluid (TIF) that represents TME from different types of tumors in mice and measured ammonia concentration in comparison to subcutaneous fluid (SCF) (Fig. 2d). In these experiments, both immunocompetent BALB/c mice and immunocompromised SCID or NSG mice were used for inoculation of murine and human cell lines, respectively. In both syngeneic and xenograft models, we found that ammonia accumulated in TME in all types of studied tumor models including lymphoma (Raji, Fig. 2e), multiple myeloma (MM.1s, Fig. 2f), and breast cancer tumors (EMT6, E0771, 4T1, MCF-7, MDA-MB 231 Fig. 2g). The concentrations of ammonia in TIFs varied between different tumors from 0.5 mM to 5 mM, which remains consistent with the already published data, and were in general even higher than those measured in cancer-conditioned medium *in vitro*.

### Ammonia suppresses NK cells

To determine whether ammonia may indeed regulate the activity of NK cells, we tested the influence of exogenously added ammonia in a range of concentrations corresponding to those observed *in vivo* in TIF on NK cells. Ammonia did not affect the viability of NK cells (Fig. 3a), even in long-term settings (Supplementary Fig. 3a). Nonetheless, we found that ammonia accumulated within NK cells (Supplementary Fig. 3b), similarly to what was demonstrated in T cells ^16^.

**Fig. 3.**
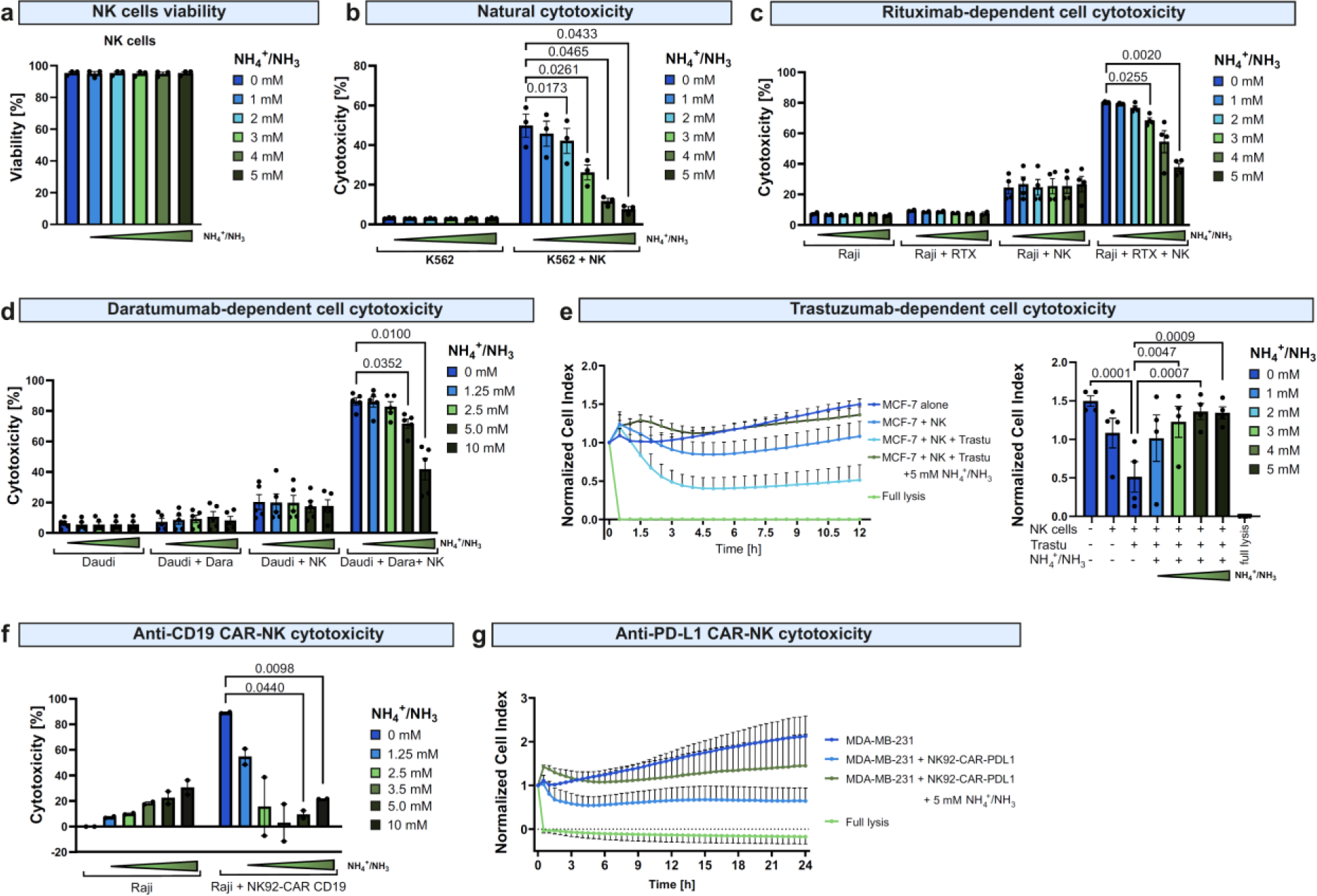
Ammonia inhibits natural cytotoxicity and ADCC of NK cells and NK-CAR cells. **a**, Viability of NK cells incubated with different concentrations of ammonia (ammonium chloride) for 4 hours was assessed using propidium iodide staining and flow cytometry (n=3). **b**, Natural cytotoxicity of NK cells against K562 cells in the presence of different concentrations of ammonia (ammonium chloride) (n=3). K562 cells were stained with CFSE and incubated with NK cells in different concentrations of ammonia. **c**, RTX-dependent cell cytotoxicity of NK cells against Raji cells in the presence of different concentrations of ammonia (ammonium chloride) (n=4). Raji cells were stained with CFSE and incubated with NK cells and 100 µg/ml RTX in different concentrations of ammonia. **d**, Dara-dependent cell cytotoxicity of NK cells against Daudi cells in the presence of different concentrations of ammonia (ammonium chloride) (n=5). Daudi cells were stained CFSE and incubated with NK cells and 1 µg/ml Dara in different concentrations of ammonia. In **b-d** cytotoxicity was assessed after 4 hours using flow cytometry and plotted as percentage of propidium iodide-positive CFSE-positive target tumor cells. **e**, Trastuzumab-dependent cell cytotoxicity of NK cells against MCF-7 cells in the presence of different concentrations of ammonia (ammonium chloride) (n=4). Cytotoxicity was assessed using Real-Time Cytotoxicity Assay (RTCA) for 12 hours. Right panel presents normalized cell index at 12-hour time point. **f**, CD19 CAR-NK cells cytotoxicity against Raji cells in the presence of different concentrations of ammonia (ammonium chloride) (n=2). Cytotoxicity was determined after 18 h in a luciferase-based killing assay with Raji stably expressing luciferase as target cells. **g**, PD-L1 CAR-NK cells cytotoxicity against MDA-MB-231 cells in the presence of 5mM ammonium chloride (n=3). Cytotoxicity was assessed using RTCA for 24 hours. *P* values were calculated using two-way ANOVA with Tukey’s post hoc test. Data show individual values and means ± SEM. *n* values are the numbers of biological replicates in *in vitro* experiments. The source data underlying **a-g** are provided as a Supplementary Data file.

Notably, ammonia inhibited the natural cytotoxicity of NK cells in a dose-dependent manner (Fig. 3b). In concentrations observed in most cancer-conditioned media and TIFs (2-3 mM), ammonia substantially suppressed NK cell cytotoxicity, while at higher (4-5 mM) concentrations that were still within a range of *in vivo* measurements it completely inhibited NK cell killing activity. Moreover, the suppression of NK cell cytotoxicity was even more pronounced when NK cells were preincubated with ammonia for up to 48 hours before contact with target cells (Supplementary Fig. 3c).

Using RTX and daratumumab we demonstrated that ammonia suppressed ADCC against Raji (Fig. 3c) and Daudi cells (Fig. 3d), reproducing the effects of a cancer-conditioned medium (Fig. 1e). Moreover, ammonia inhibited trastuzumab-dependent ADCC against breast cancer MCF-7 cells, as assessed by real-time cell analysis (RTCA) assay (Fig. 3e). In addition to the engagement of NK cells by monoclonal antibodies, NK cells can be redirected to the tumor cells by introduction of tumor-targeting chimeric antigen receptors (CAR). Using NK-92 cell line transduced with CD19 (FMC-63 scFv) or PD-L1 (atezolizumab-based) CAR, we found that ammonia suppressed cytotoxicity of NK-92 CAR cells against Raji (Fig. 3f) and MDA-MB-231 breast cancer cells (Fig. 3g), respectively. Altogether, this data demonstrates that ammonia potently suppresses the antitumor activity of NK cells and NK cell-based therapeutic strategies.

To further determine the effects of ammonia on the antitumor activity of NK cells *in vivo*, we used BALB/c SCID mice characterized by an absence of functional T cells and B cells, that were inoculated subcutaneously with Raji cells. Mice were treated systemically with 10 mg/kg RTX and 50 mM of ammonium chloride was administered intratumorally (Supplementary Fig. 4a). Only a trend toward shorter survival (Supplementary Fig. 4b) and a higher tumor volume (Supplementary Fig. 4c) of mice that were injected intratumorally with ammonia and RTX as compared to the RTX-only group was observed and it did not reach statistical significance. When RTX and ammonia were administered for a longer time (Supplementary Fig. 4d), we observed similar results (Supplementary Fig. 4e). These results motivated us to explore this issue further. According to the previous study, ammonia administered systemically as a bolus of 9 mmol/kg ammonium chloride is rapidly recycled in TME into amino acids by cancer cells ^10^. We found that after local administration of 50 mM of ammonium chloride directly into tumor masses (Supplementary Fig. 5a), it is rapidly utilized or excreted from TME since no significant changes in the ammonia concentration were detected in the TIFs of ammonia-injected tumors after 0.5 or 4 hours (Supplementary Fig. 5b). Finally, we found that even administration of a supraphysiological dose of ammonium chloride (100 mM) into TME did not change ammonia concentration in the TIFs after as early as 30 min (Supplementary Fig. 5c). Nonetheless, based on *in vitro* data and *ex vivo* ammonia measurements in TIF isolated from tumor masses, we sought to determine the mechanism of the suppressive effects of ammonia on NK cell activity.

### Ammonia decreases the level of perforin in NK cells

The process of killing target cells by NK cells consists of several steps, which include NK cell activation, conjugation with a target cell, degranulation, cytokine production, and detachment from target cells ^21^. We found that ammonia did not affect the surface levels of NK cell activating receptors NKG2D, NKp30, NKp44, and NKp46 (Fig. 4a). Moreover, there was no impairment of the formation of conjugates of NK and target cells in the presence of ammonia (Fig. 4b). After formation of conjugates, ammonia did not influence the percentage of degranulating NK cells (Fig. 4c). Unexpectedly, high ammonia concentration increased the percentage of TNF-α- and IFN-γ-expressing NK cells (Fig. 4d). Moreover, we observed delayed detachment of NK cells from target cells in a high ammonia concentration (Fig. 4e). These findings suggested that in the presence of ammonia, NK cells are still fully functional to attach to target cells and release the content of secretory lysosomes (degranulate). Nonetheless, they are unable to execute target cell killing which is a signal for the disengagement from target cells ^22^. Therefore, we determined the levels of granzyme B and perforin which are two major cytolytic proteins within secretory lysosomes ^23^. While the level of granzyme B was unaffected even in high concentrations of ammonia (Fig. 4f), we found that the level of perforin in NK cells was significantly decreased by ammonia in a dose-dependent manner (Fig. 4g).

**Fig. 4.**
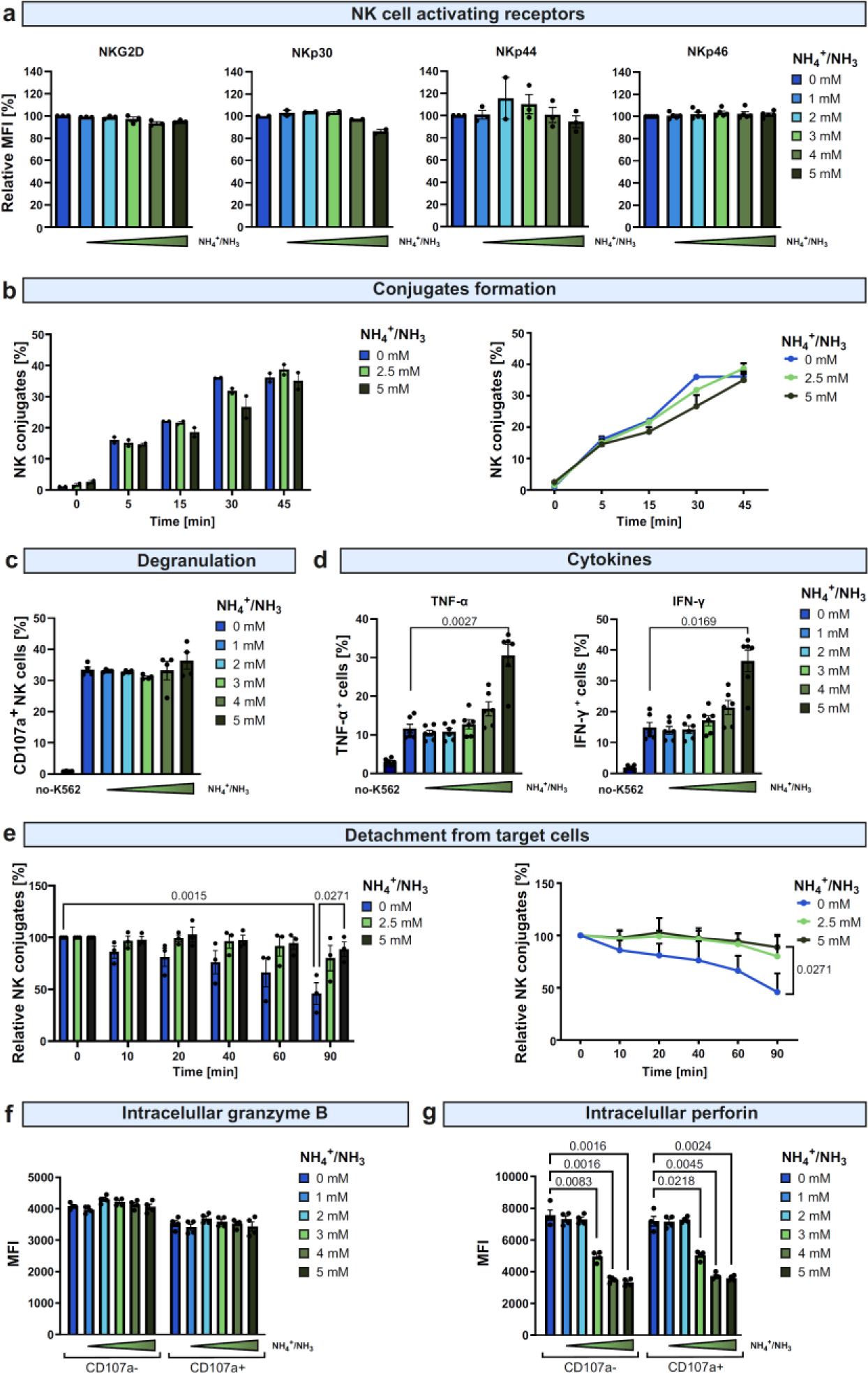
Ammonia decreases the level of intracellular perforin. **a**, The surface level of NK cell activation receptors (NKG2D, NKp30, NKp44, NKp46) after incubation with different concentrations of ammonia (ammonium chloride). The levels of receptors were assessed after 4 hours using flow cytometry. Mean fluorescence intensity (MFI) was normalized to control (medium without exogenously added ammonium). **b,** Formation of conjugates of NK and target cells in different concentrations of ammonia (ammonium chloride) (n=3). NK cells were stained with Cell Trace Deep Red (CTDR) and target cells (K562 cells) were stained with CFSE. Cells were mixed and incubated with different concentrations of ammonia at 37°C for 30 minutes. After this time, 4% paraformaldehyde (PFA) was added followed by 10 minute incubation and wash. The percentage of conjugates defined as CTDR^+^CFSE^+^ double-positive events was determined using flow cytometry. Graph shows representative experiment from one donor. **c,** Degranulation of NK cells in response to stimulation with target cells (K562) in different concentrations of ammonia (ammonium chloride) (n=4). The percentage of degranulating NK cells (CD107a^+^) was assessed after 4 hours using flow cytometry. **d,** the level of cytokines expressed in NK cells in response to stimulation with target cells (K562) in different concentrations of ammonia (ammonium chloride) (n=6). The percentage of tumor necrosis factor alpha (TNF-α)-positive and interferon gamma (IFN-γ)-positive NK cells was assessed after 4 hours using flow cytometry. **e**, Detachment of NK cells from target cells in different concentrations of ammonia (ammonium chloride) (n=3). NK cells were stained with CTDR and target cells (K562 cells) were stained with CFSE. Cells were mixed and incubated in different concentrations of ammonia. After 0-45 minutes, 4% PFA was added followed by 10-minute incubation and wash. The percentage of conjugates defined as CTDR^+^CFSE^+^ double-positive events was determined using flow cytometry. **f**, the level of granzyme B in NK cells incubated in different concentrations of ammonia (ammonium chloride) (n=4). The level of granzyme B in CD107a^-^ and CD107a^+^ NK cells was assessed by intracellular staining using flow cytometry. **g,** The level of perforin in NK cells incubated in different concentrations of ammonia (ammonium chloride) (n=4). The level of total perforin in CD107a^-^ and CD107a^+^ NK cells was assessed by intracellular staining (B-D48 antibody) using flow cytometry. *P* values were calculated using two-way ANOVA with Tukey’s post hoc test. Data show means ± SEM. *n* values are the numbers of biological replicates in *in vitro* experiments. The source data underlying **a-g** are provided as a Supplementary Data file.

### Ammonia decreases the amount of mature perforin

We further confirmed the decrease of perforin level in NK cells by ammonia using a western blot method (Fig. 5a). Similar downregulation of perforin was observed when NK cells were incubated in a cancer-conditioned medium (Supplementary Fig. 6a). Notably, ammonia not only decreased intracellular perforin in NK cells but also decreased the total amount of perforin that was secreted by the NK cells in response to cancer cell recognition (Fig. 5b). We observed a similar trend toward decreased extracellular perforin level when NK cells were incubated with target cells in cancer-conditioned media (Supplementary Fig. 6b-d). Notably, the downregulation of perforin by low doses of ammonia was reversible since after 16 hours of ammonia washout the amount of perforin reversed back to the control levels (Fig. 5c). These findings suggest that the decrease of perforin upon contact with ammonia is a relatively fast process, most probably related to its maturation and/or degradation.

**Fig. 5.**
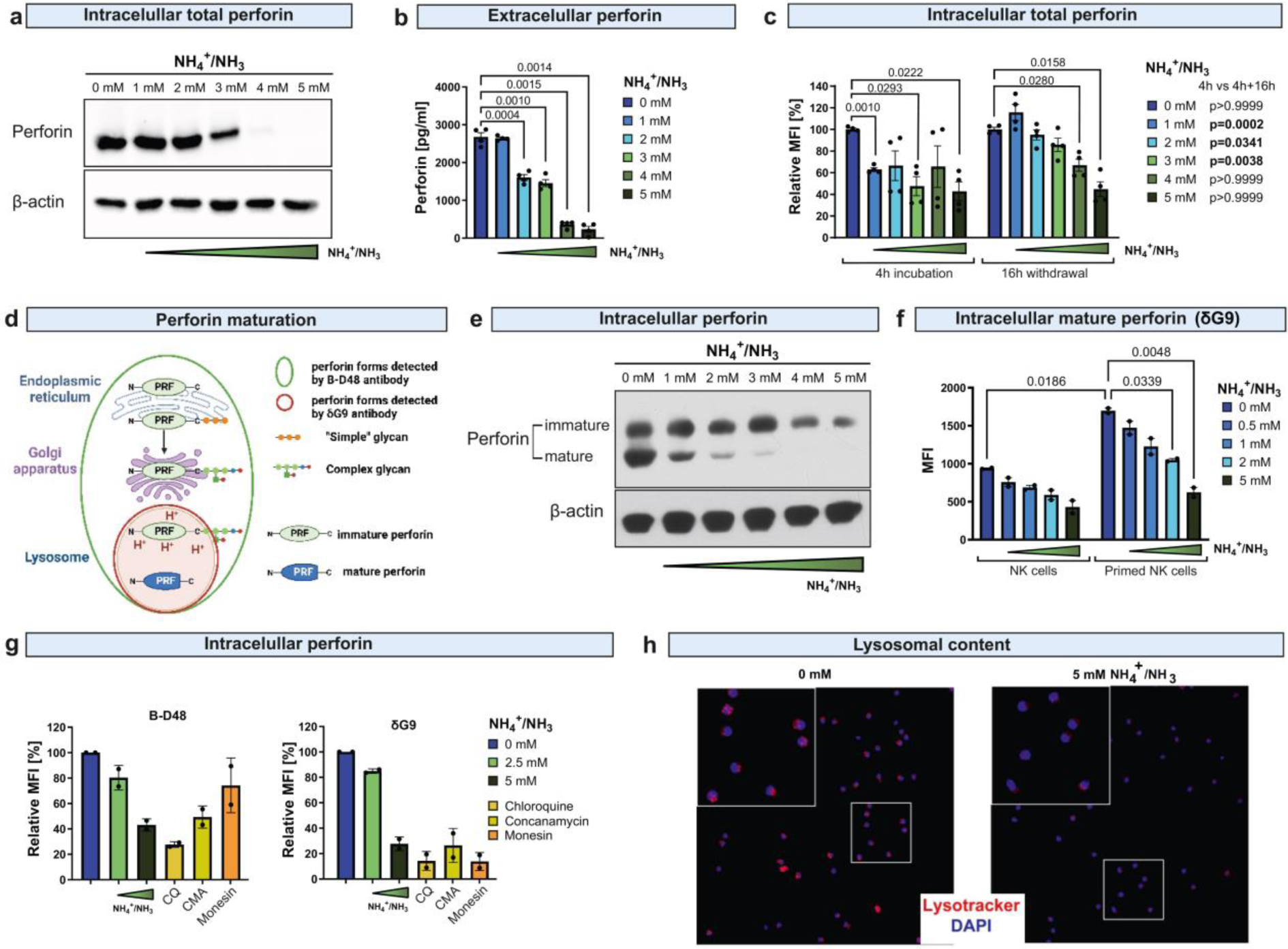
Ammonia impairs perforin maturation. **a**, The level of total perforin in NK cells incubated with ammonia determined by western blot methods using anti-perforin antibody (B-D48 clone) (n=3). β-actin presented as a loading control. Representative blot from one donor. **b,** The concentration of extracellular perforin secreted by NK cells in response to contact with target cells (K562) in different concentrations of ammonia (ammonium chloride) (n=4). **c,** The level of perforin detected in NK cells incubated with ammonia (ammonium chloride) determined by intracellular staining using anti-perforin antibody (δG9 clone) and flow cytometry (n=4). In some groups, cells were washed and incubated in a medium without ammonia as indicated in the figure. **d**, A schematic representation of perforin maturation. Green and red circle shows forms of perforin recognized by B-D48 and δG9 antibodies, respectively **e,** The level of perforin in NK cells incubated with ammonia (ammonium bicarbonate) determined by western blot methods using anti-perforin antibody (Pf-344 clone) (n=3). Two forms of perforin, immature (70 kDa) and mature (60 kDa) were detected. **f**, The level of perforin detected in NK cells incubated with ammonia (ammonium chloride) determined by intracellular staining using anti-perforin antibody (δG9 clone) and flow cytometry (n=2). NK cells were primed with IL-2 (200 U/ml) and IL-15 (10 ng/ml) for 24 hours before the experiment. **g**, The level of perforin detected in NK cells incubated with ammonia (ammonium chloride) and other lysosomotropic agents (chloroquine (CQ), concanamycin (CMA), and monesin) incubated for 4 hours determined by intracellular staining using antibody detecting total perforin (B-D48 clone) and lysosomal perforin (δG9 clone) (n=2). **h,** The lysosomal content of NK cells incubated with 0 or 5mM ammonia (ammonium chloride) for 12 hours. Acidic organelles were stained using a LysoTracker and live imaged using Opera Phenix cell microscopy (n=4). Data show representative images, the effect of ammonia over the increasing concentration range is shown in supplement. The source data underlying **a-h** are provided as a Supplementary Data file.

Perforin is synthesized as a 70 kDa inactive precursor that undergoes several posttranslational modifications including proteolytic cleavage in an acidic compartment yielding a 60 kDa mature perforin which is stored in the lysosomes until secretion in response to stimuli (Fig. 5d). Thus, it is possible to distinguish immature and mature perforin based on their molecular size ^24–26^. Indeed, we detected both forms of perforin in primary NK cells using Pf-344 antibody that binds a linear epitope within EGF-like domain ^27^. Notably, ammonia decreased mature perforin level even in as low concentration as 1 mM (Fig. 5e). Mature perforin completely disappeared in the high (4-5 mM) ammonia concentrations. Further, we confirmed by flow cytometry the reduction of mature perforin amount using δG9 antibody (Fig. 5f) that recognizes pH-sensitive motif in granule-associated mature perforin (Fig. 5d, ^28^). Mature perforin was severely decreased in NK cells by ammonia which was even more pronounced in NK cells primed with IL-2 and IL-15 that potently expressed perforin (Fig. 5f).

The most important stages of perforin maturation in NK cells take part in specific organelles that provide necessary acidic pH, including secretory lysosomes (Fig. 5d, ^29^). Since it was reported that ammonia is a lysosomotropic agent ^30^, we hypothesized that the decreased amount of perforin may be caused by increased pH in the lysosome. We found that other lysosomotropic agents, including chloroquine, concanamycin, and monesin, exerted similar to ammonia effects on perforin with most pronounced changes in mature perforin detected by a δG9 antibody (Fig. 5g). To determine whether changes in acidic compartment mediate effects of ammonia on perforin, we utilize a LysoTracker probe which selectively accumulates in acidic organelles. Indeed, we demonstrated that ammonia decreased acidic compartment in NK cells (Fig. 5h, Supplementary Fig. 7a). Then, we analyzed the CD63 (lysosomal-associated membrane protein 3, LAMP3) and LAMP1 (CD107a), markers of secretory lysosomes ^31^ in NK cells incubated with different doses of ammonia (Supplementary Fig. 7b). We observed that ammonia increased the number of LAMP1^+^ vesicles (Supplementary Fig. 7c) but decreased their area (Supplementary Fig. 7d) and mean LAMP1 level in NK cells (Supplementary Fig. 7e). Ammonia had no effect on CD63^+^ vesicles. It suggests that ammonia selectively upregulates pH in acidic organelles. Collectively, we demonstrated that ammonia decreases mature perforin level and that this effect is related to the changes in the acidic compartment of NK cells.

### Ammonia inhibits NK cell serial killing

Finally, we sought to determine the consequences of ammonia-impaired effector functions of NK cells. Decreased cytotoxicity of NK cells was associated with the increased percentage of NK cells that degranulated more than once (serial degranulation) (Fig. 6a), which may be caused by the lack of signals promoting NK cell detachment and a second attempt to kill the target cell. Ammonia significantly increased the membrane expression of CD56 on both resting and activated by targets NK cells (Fig. 6b). Moreover, it increased the levels of both Fas (CD95, Fig. 6c) and FasL (CD178, Fig. 6d) on NK cells, suggesting a switch to death receptor-mediated cytotoxicity.

**Fig. 6.**
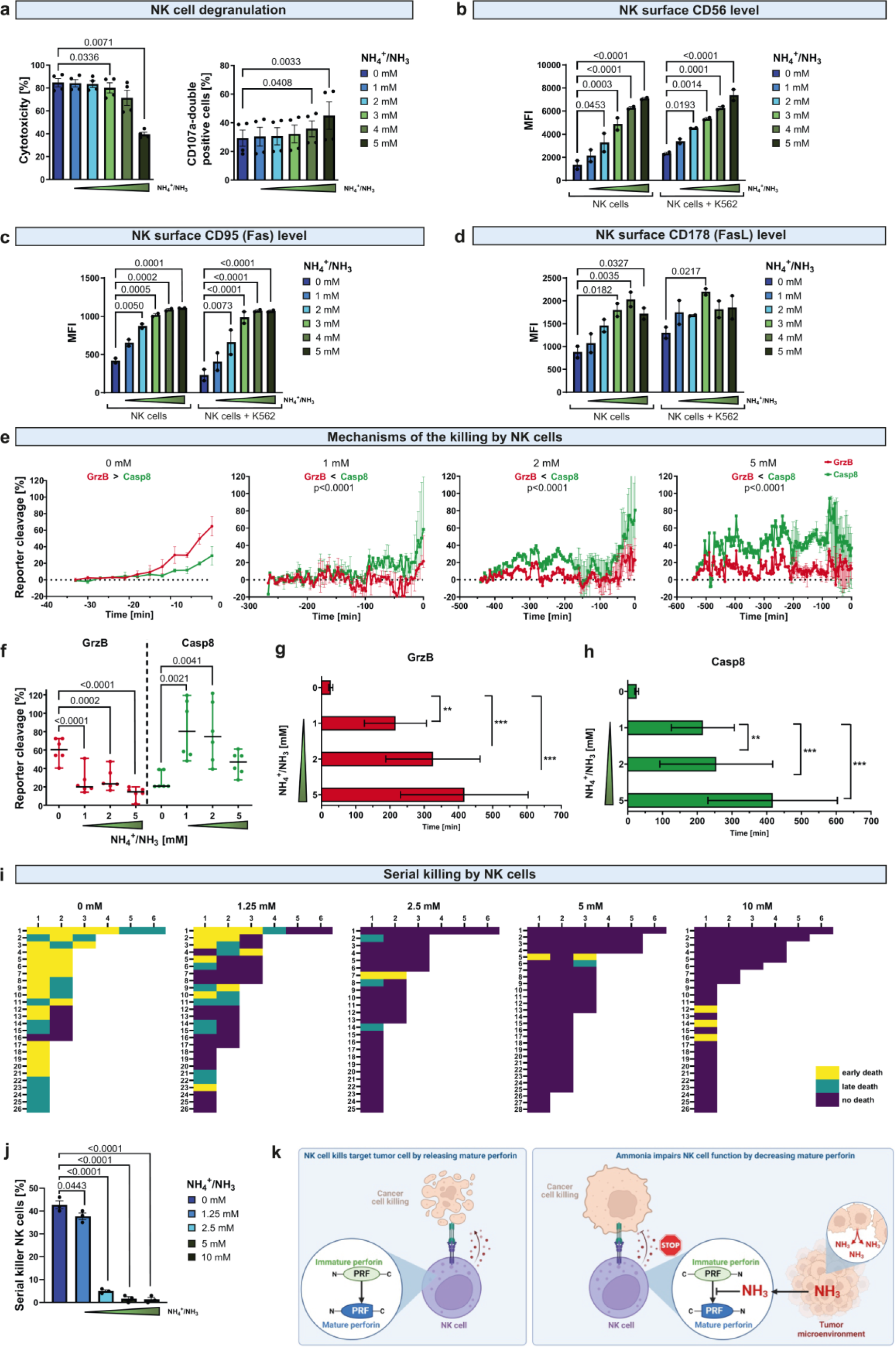
Ammonia inhibits NK cell serial killing. **a**, The percentage of NK cells that underwent serial degranulation in response to stimulation with target cells (K562) in different concentrations of ammonia (ammonium chloride) (n=4). Cells were stained with anti-CD107a after 2 hours of incubation, and then after additional 2h were stained with anti-CD107a antibody conjugated with a different fluorochrome and analyzed using flow cytometry. NK cells that underwent serial degranulation were defined as CD107a-double positive NK cells. **b,** The level of CD56 in NK cells incubated with ammonia (ammonium chloride) with or without target cells determined by surface staining and flow cytometry (n=2). **c,** The level of CD95 (Fas) in NK cells incubated with ammonia (ammonium chloride) with or without target cells determined by surface staining and flow cytometry (n=2).**d,** The level of CD178 (FasL) in NK cells incubated with ammonia (ammonium chloride) with or without target cells determined by surface staining and flow cytometry (n=2). **e-k,** HeLa cells were transfected with NES-ELQTD-GFP-T2A-NES-VGPD-mCherry and CD48. In these cells, NES-RIEADS-mCherry (activation of GrzB) and NES-VGPD-mGFP (activation of caspase 8) reporter cleavage can be detected by the appearance of fluorescence inside the nucleus. GrzB cleaves the reporter, resulting in an increase of red fluorescent signal in the nucleus, while Casp8 reporter is specifically activated by death receptor–mediated apoptosis and results in an increase of green fluorescence in the nucleus. Confocal time-lapse microscopy started immediately after NK cell exposure. **e,** Activation of granzyme B (GrzB) and caspase 8 (Casp) reporters in target HeLa cells incubated with NK cells in different concentrations of ammonia (ammonium chloride). Time point 0 sets to the time of cell death (n=15). Images were acquired every 3 min. **f,** Activation of granzyme B (GrzB) and caspase 8 (Casp) reporters in target HeLa cells incubated with NK cells in different concentrations of ammonia (ammonium chloride) at the time of cell death (n=6). **g,** Time required to kill a reported-expressing cell by NK cell in different concentrations of ammonia (ammonium chloride) by activation of granzyme B (n=15). SYTOX^TM^ Blue dye was used to assess the viability of the target cell. Early or late cell death was evaluated based on the kinetics of the target cell nucleus’ staining after the contact with NK cells. **h,** Time required to kill a reported-expressing cell by NK cell in different concentrations of ammonia (ammonium chloride) by activation of caspase 8 (n=15). **i**, Diagram displays target cell death in single killing events. Each row displays the killing sequence of one individual NK cell imaged using time-lapse microscopy (controls, n=26; 1.25 mM, n=26; 2.5 mM, n=25; 5 mM, n=28; 10 mM, n=27). **j,** Percentage of serial killers among NK cells in different concentrations of ammonia (ammonium chloride) (controls, n=26; 1.25 mM, n=26; 2.5 mM, n=26; 5 mM, n=28; 10 mM, n=27). *P* values were calculated using two-way ANOVA with Tukey’s post hoc test. Data show means ± SEM. *n* values are the numbers of biological replicates in *in vitro* experiments. **k**, A schematic representation of the main findings. The source data underlying **a-j** are provided as a Supplementary Data file.

To verify whether ammonia inhibits granzyme-dependent killing and leads to a compensatory upregulation of death receptor-mediated cytotoxicity, we used a Hela cell line that expresses two reporters enabling quantification of granzyme B and caspase 8 activity in living cells by fluorescence measuring ^32^. This method allows live tracking of target killing by NK cells and distinguishes between perforin/granzyme B-mediated death and death receptor-mediated cell killing ^33^. Live cell imaging of control NK cells incubated with reporters-expressing cancer cells revealed that the dominant mechanism of killing by NK cells depends on perforin/granzyme B. However, in the presence of ammonia, this mechanism was severely impaired even in low concentrations of ammonia (Fig. 6e). Ammonia induced a switch from perforin/granzyme B to death receptor-mediated cytotoxicity as revealed by the increased cleavage of caspase 8 reporter (Fig. 6f). Notably, live cell imaging revealed that even in low ammonia concentration NK cells needed significantly more time to kill the target cell. In 2 mM of ammonia, the time from the contact with the target cell to successful killing was prolonged about 10 times, from 27 min for granzyme B and 25.5 min for caspase 8 to 254.5 min and 325.25 min, respectively (Fig. 6g).

An important feature of NK cells is the ability to detach from the target cell followed by binding and killing other cells in a process called serial killing ^22^. However, this process is largely limited by the availability of lytic granules containing perforin and granzyme B ^34,35^. To investigate whether ammonia affects serial killing, we tracked the killing sequence of NK cells incubated with cancer cells in individual microwells. We found that ammonia significantly decreased the percentage of serial killers among NK cells (Fig. 6i). Each NK cell was able to kill on average 1.5 target cells (Fig. 6j). However, in the presence of ammonia average number of cells killed by one NK cell substantially decreased leading to the profound reduction of the population of NK cells that were serial killers (Fig. 6k). In the presence of ammonia most of the NK cell-target cancer cell interactions were unsuccessful and did not lead to the death of target cells. These results revealed that ammonia, by impairing perforin-mediated cell killing, substantially prolongs the time needed for NK cells to kill cancer cells, inhibits serial killing, induces a switch toward death receptor-mediated cytotoxicity, and significantly reduces the effectiveness of cancer cell killing after recognition by NK cells.

## DISCUSSION

NK cell-based immunotherapies are currently extensively studied in preclinical and clinical studies as a promising alternative to T cells ^36–38^. This field currently involves CAR-based therapies, monoclonal antibodies with modified Fc to enhance ADCC as well as bi-specific and tri-specific NK cell-engaging antibodies. Nonetheless, TME remains one of the main obstacles to the effectiveness of NK cell-based therapy ^4^. Here, we identified that microenvironmental ammonia is a metabolic immune checkpoint for NK cells. We demonstrated that the cancer-conditioned medium suppressed the antitumor activity of NK cells. Ammonia accumulated in a conditioned medium and TME reaching ∼2-5 mM, which is similar to concentrations reported in previous studies ^10,20,39^. In these concentrations, ammonia inhibited the cytotoxicity of NK cells and decreased total and mature perforin levels leading to NK cell dysfunction (Fig. 6l).

Recently ammonia arose as an important player in cancer. While systemic hyperammonemia in cancer patients is rather rare and generally does not exceed micromolar concentrations ^40^, ammonia accumulates in TME reaching millimolar concentrations. It is generated during a glutamine breakdown into glutamate by glutaminase (GLS) ^41^. In cancer, ammonia is recycled and serves as a source of nitrogen to build amino acids promoting the growth of tumor biomass as well as supports autophagy and protects cells from TNF-α-induced cell death ^10,20,42^. Conversely, in high concentrations, ammonia suppresses polyamine biosynthesis and inhibits cancer cell proliferation ^43^. Ammonia can be utilized by three enzymes, carbamoyl phosphate synthase-1 (CPS1), glutamine synthetase (GS), and glutamate dehydrogenase (GDH) which expression is regulated by oncogenes and growth signaling pathways ^44^. Low expression of ammonia-detoxifying CPS1 is associated with worse overall survival of hepatocellular cancer patients ^45^. A high ammonia gene expression signature which includes low expression of ammonia-detoxifying enzymes and high expression of ammonia-producing enzymes is associated with decreased survival of patients with gastrointestinal carcinomas ^16^. Notably, other types of cells in TME also contribute to the nitrogen metabolism and thus production of ammonia ^41,44^.

Furthermore, ammonia promotes cancer progression indirectly by suppressing antitumor immune response. A recent study demonstrated that ammonia inhibits T cell-dependent antitumor immunity and enhances their exhaustion ^16^. Gene signature associated with high ammonia was a predictor of worse survival following immune checkpoint inhibitors ^16^. Moreover, ammonia inhibits antigen presentation by macrophages, phagocytosis by neutrophils, and impairs functions of dendritic cells ^11–15^. Here, we demonstrated that also NK cells are impaired by ammonia, thus suggesting its wide suppressive effects on immune response.

It is well established that cancer cells and tumor-promoting immune cells suppress the antitumor activity of NK cells. Different cytokines including TGF-β, IL-6, and IL-10, as well as cancer-associated metabolites, including tryptophan metabolites, lactate, and prostaglandin E2 inhibit NK cell functions at various levels ^4^. These factors regulate the expression of chemokine receptors, activating and inhibitory receptors, cytokines, and molecules of the lytic granule, granzymes and perforin.

Perforin has a primary role in the NK cell-mediated control of tumor initiation, growth, and metastasis ^46–49^. It is necessary for granule-dependent target cell death since it enables granzyme-induced apoptosis ^29^. Without perforin, NK cells are unable to perform granzyme B-mediated serial killing and only kill via death receptors which significantly prolongs the time required to kill cancer cells ^33,50^. Here, we observed that ammonia significantly impairs the antitumor activity of NK cells by decreasing perforin. Perforin is synthesized as a 70 kDa inactive precursor that is subjected to different modifications to yield a mature 60 kDa active form ^25^. Trafficking of perforin from the endoplasmic reticulum, through Golgi to the lysosomal compartment is accompanied by the proteolytic cleavage and LAMP1/CD107a ^51^. NK cells with LAMP1 knockdown have impaired perforin recruitment to lytic granules which leads to the inhibition of their cytotoxicity ^51^. The activity of mature perforin is regulated by two main factors, pH and calcium ions ^29^. Moreover, mature perforin is stored in lysosomal granules (secretory lysosomes) whose low pH prevents oligomerization protecting cytotoxic lymphocytes ^52^. In secretory lysosomes, the lytic activity of perforin is prevented by chaperon protein calreticulin and proteoglycans ^53,54^. This association of perforin with granule proteins is pH-dependent and increased pH leads to the dissociation of perforin from the proteoglycans ^54,55^. Here, we demonstrated that ammonia increases pH in the lysosomes and decreases amount of mature perforin. Previous studies demonstrated that alkalization of acidic compartment impairs perforin maturation by inhibiting proteolytic cleavage ^25^ as well as leads to its inactivation and proteolytic degradation ^56^. In our study, we observed the rapid disappearance of mature perforin in low concentration of ammonia and decrease of immature perforin upon treatment with 4-5 mM ammonia (Fig. 5e). It may suggest that both impairment of perforin maturation and its proteolytic degradation play a role in the effects of ammonia on NK cells.

In the presence of ammonia, NK cells compensatory upregulated the expression of death receptors (Fig. 6c-d). Nonetheless, they still had impaired cytotoxic functions as they needed significantly more time to kill target cells (Fig. 6g) and the majority of their contacts with target cells did not lead to successful killing (Fig. 6i). Previous studies demonstrated that defective killing results in an enhanced cytokine secretion which is caused by failed detachment from target cells ^22,50^. Indeed, we observed that ammonia increased levels of IFN-γ and TNF-α in NK cells (Fig. 4d). Such delayed detachment was observed in human NK cells with genetic perforin knockout ^22^. Accordingly, we observed that ammonia delayed the disengagement of NK cells from their targets (Fig. 4e). Notably, previous studies revealed that ammonia does not affect target cell sensitivity to killing by purified granzyme B and perforin ^57^, further confirming selective impairment of NK cells cytotoxic machinery by ammonia.

Importantly, different components of TME including hypoxia, PGE2, lactate, and kynurenines decrease perforin levels in NK cells ^58–61^. However, the molecular mechanisms of these effects remain unknown. We observed that ammonia even in low concentration potently decreased the amount of mature perforin. It is known that genetic mutations in the perforin gene may result in the aberrant maturation that is observed in hemophagocytic lymphohistiocytosis ^24^. Here, we identified that ammonia decreases the amount of mature perforin by increasing the pH of the acidic compartment which results in a decreased level of total perforin and severely impairs perforin/granzyme B-mediated killing.

Ammonia (NH_3_) is neutral and rapidly passes across biological membranes. In contrast, ammonium ion (NH_4_^+^) cannot diffuse through membranes and accumulates in acidic organelles, including lysosomes. A recent study identified SLC12A9 as a lysosome-detoxifying ammonium-chloride co-transporter ^62^. However, its expression in immune cells remains unknown. Moreover, while hepatocytes or tumor cells can metabolize ammonia, immune cells like T cells, macrophages, and NK cells lack this feature (Supplementary Fig. 3b, ^16^), which may explain its immunosuppressive effects. Recent studies demonstrated that CD8^+^ T effector (T_eff_) and memory (T_mem_) cells can detoxify ammonia ^17,18^. However, whether some populations of NK cells possess that feature remains unknown. Still, metabolic impairment is one of the key factors driving NK cells dysfunction in tumors and reprogramming of the metabolic fitness can strengthen their antitumor activity ^63^.

Therapeutic strategies that decrease ammonia in TME may restore NK cell activity against the tumors. It is supported by the fact that the effects of low ammonia concentrations on the level of perforin are fully reversible (Fig. 5c). Previous study demonstrated that neutralization of ammonia reduced colorectal cancer tumor growth and enhanced T cell response to immune checkpoint inhibitors ^16^. Moreover, there are other strategies to modulate ammonia levels in TME. A non-pathogenic strain of *E. coli* was engineered to use ammonia for continuous arginine synthesis ^64,65^. This strain recycles ammonia and elevates the arginine amount necessary for T cells in TME leading to activation of T cell response, inhibition of tumor growth, and prolonged survival of tumor-bearing mice ^64,66^. On the other side, a high ammonia level may be used as a marker of TME to selectively deliver different cytotoxic compounds to tumors ^39^.

Together, our results for the first time reveal that ammonia is an important metabolic immune checkpoint for NK cells. It suppresses the spontaneous antitumor immune response of NK cells as well as NK cell-based therapies. Therefore, strategies to enhance ammonia detoxification may have the potential to increase the efficiency of various immunotherapies, especially those relying on the activation of NK cells.

## MATERIALS AND METHODS

### Cell culture

Cells were cultured in RPMI-1640 (Raji, Daudi, Ramos, MCF-7) (Gibco) or DMEM (E0771, EMT6) (Sigma-Aldrich) supplemented with 10% fetal bovine serum (FBS) (HyClone) and 1% penicillin/streptomycin solution (Gibco) in a humidified atmosphere containing 5% CO_2_ at 37°C. Multiple myeloma cell lines (MM1s, RPMI 8226, H929) were cultured in RPMI-1640 medium additionally supplemented with 50 mM 2-mercaptoethanol (1000x) (Gibco) and 100 mM sodium pyruvate (100x) (Gibco). NK-92 cell line was maintained in X-VIVO™ 20 medium (Lonza) supplemented with 5% human serum (Sigma Aldrich). Cell media was changed every 48 hours. For 3D cell culture, 5 × 10^3^ 4T1 cells were seeded in a 48-well plate on an LDEV-free MatriGel (Corning) in RPMI supplemented with 5% FBS, 1% penicillin/streptomycin, and 2% MatriGel. Cells were cultured for 14 days. Conditioned medium was collected following 48 hours of incubation. Amicon Ultra-0.5 Centrifugal Filter Unit Ultracel-3 was used according to the manufacturer’s protocols to separate a small-molecule fraction (< 3 kDa) and large-molecule fraction (> 3 kDa) from the conditioned medium. Cells were regularly tested for Mycoplasma contamination using the PCR method.

### NK cell isolation

Peripheral Blood Mononuclear Cells (PBMCs) were isolated from buffy coats of healthy donors by gradient centrifugation using Ficoll (Sigma Aldrich). Buffy coats were purchased from the Regional Blood Centre in Warsaw, the procedure was approved by the Local Bioethics Committee (approval number: AKBE/61/2021). Primary NK cells were isolated from PBMCs by immunomagnetic negative selection using EasySep™ Human NK Cell Enrichment Kit (STEMCELL Technologies Canada, Inc.) according to the manufacturer’s protocols. NK cells were cultured in full RPMI-1640 medium unless otherwise described for specific experimental procedures.

### Animal studies

All *in vivo* experiments were performed with 8–12-week old female mice obtained from Charles River Laboratories, which were bred at the animal facility of the Department of Immunology, Medical University of Warsaw. All experiments were performed in accordance with the guidelines and approved by The Second Local Ethics Committee for the Animal Experimentation, Warsaw University of Life Sciences (number: WAW2/005/2021, WAW2/074/2021, WAW/175/2022).

### *In vivo* tumor models

Experiments were performed on NSG (ang. NOD scid gamma mouse), SCID, Balb/c or C57BL/6 female mice. NSG mice were inoculated subcutaneously with human cell lines: 2 × 10^6^ Raji, MDA-MB-231 or 3 × 10^6^ MCF-7 cells. In the case of the experiments involving MCF-7 cells the slow-release pellets containing 17β-estradiol (Innovative Research of America, USA), were implanted subcutaneously four days before tumor cells inoculation. SCID mice were inoculated subcutaneously with 3 × 10^6^ MM.1s cells. BALB/c mice were inoculated into the left mammary fat pad with murine cell lines: 0.37 × 10^6^ 4T1 cells or subcutaneously with 0.37 × 10^6^ EMT6 cells. C57BL/6 mice were inoculated subcutaneously with 0.37 × 10^6^ E0771 cells. Tumor volume was calculated according to the formula mm^3^= (width^2^ [mm] × length [mm])/2. Mice were sacrificed when the tumor diameter reached 15 mm in at least one dimension. Tumors were used for tumor interstitial fluid (TIF) and subcutaneous fluid (SCF) isolation.

### Tumor interstitial fluid collection

Tumor interstitial fluid (TIF) was isolated from tumors not exceeding 1500 mm^3^. Mice were anesthetized by administrating 10 mg ketamine and 1.5 mg xylazine per 100 g body weight. TIF was isolated using a UF-1-2 *In Vivo* Ultrafiltration Sampling Probes (BASI, MF-7027). The probe was implanted centrally into the tumor for 2h to collect TIF. The subcutaneous fluid (SCF) of healthy tissue was extracted by a probe implanted under the skin at the opposite side of the tumor. High-molecular-weight compounds were excluded from the analytes utilizing the filtration membrane. Following ultrafiltration, approximately 10 μL of TIF and 12-18 μL of SCF was obtained. Ammonium was immediately measured using a Dimension Ammonia assay (SIEMENS).

### In vivo therapy with rituximab and intratumoral administration of ammonium chloride

Experiments were performed on BALB/c, BALB/c severe combined immunodeficiency (SCID) and NOD.Cg-*Prkdc^scid^ Il2rg^tm1Wjl^*/SzJ (NSG) mice. Mice were inoculated subcutaneously with 2 × 10^6^ Raji cells in 50% Matrigel Growth Factor Reduced (Corning, LifeSciences). Rituximab-treated mice were injected intraperitoneally with 10 mg/kg rituximab 3 times per week for 2 consecutive weeks. NH_4_Cl-treated mice were injected intratumorally with 50 mM NH_4_Cl daily for 2 consecutive weeks. Tumor volumes were measured three times a week, starting from day 7 of the experiment. Tumor volume was calculated according to the formula mm^3^= (width^2^ [mm] × length [mm])/2. Mice were sacrificed when the tumor diameter reached 15 mm in at least one dimension.

### Ammonia measurement

Ammonia concentration was measured in tumor-conditioned medium collected after 48h incubation or in TIF and SCF immediately after collection. Ammonia concentration in conditioned medium was measured using the Dimension Ammonia assay (Siemens). High-molecular-weight compounds were excluded from the analytes by a filtration membrane in the process of TIF and SCF isolation. 0.5 ml conditioned medium was centrifuged (5 min, 500g) and filtered through Amicon Ultra-0.5 Centrifugal Filter unit (Merck). Ammonia in TIFs and SCFs was then quantified using Dimension Ammonia assay (Siemens) by the Laboratory of the Central Clinical Hospital of the University Clinical Center of Warsaw, Medical University of Warsaw, Poland.

### Antibodies

Fluorophore-conjugated antibodies specific for cell-surface antigens and cytokines were as follows: anti-CD16, clone 3G8 (BioLegend); anti-CD56, clone B159 (BD Biosciences); anti-NKp30, clone p30-15 (BD Biosciences); anti-NKp44, clone p44-8 (BD Biosciences); anti-NKp46, clone 9E2 (BD Biosciences); anti-NKG2D, clone 1D11 (BD Biosciences); anti-CD95, clone DX2 (BioLegend); anti-CD178, clone NOK-1 (BD Biosciences); anti-CD107a, clone H4A3 (BD Biosciences); anti-CD107a, clone H4A3 (BD Biosciences); anti-perforin, clone δG9 (BioLegend and BD Biosciences); anti-perforin, clone D48 (BioLegend); anti-granzyme B, clone GB11 (BD Biosciences); anti-IFN-γ, clone 25723.11 (BD Biosciences); anti-TNF-α, clone MAb11 (BD Biosciences). Antibodies were used in a concentration recommended by a manufacturer.

### Flow cytometry analysis

Flow cytometry was performed on FACSCanto II (BD Biosciences), Fortessa X20 (BD Biosciences) or FACSCalibur (BD Biosciences) operated by FACSDiva software. For data analysis FlowJo v10.6.1 software (Tree Star) or BD FACSDiva software (BD Biosciences) were used. For cell surface staining, cells were stained with Zombie NIR™ or Zombie Aqua™ Fixable Viability Kit (BioLegend) for 15 minutes in RT, washed with FACS buffer (PBS; 1% BSA, 0.01% sodium azide), blocked on ice with 5% normal rat serum in FACS buffer, and then incubated for 30 min on ice with fluorochrome-labeled antibodies listed above. After washing in FACS buffer, cells were immediately analyzed.

### Intracellular staining

Cells were seeded at density 0.2 × 10^6^ NK and 0.1 × 10^6^ K562 cells (E:T ratio 2:1) per well of U-bottom plate and coincubated for 4h at 37°C in the presence of NH_4_Cl and 1 µl Golgi PLUG (BD Biosciences). After that time cells were washed three times with 200 µl PBS and centrifuged (5 min, 500g). Cells were stained with viability dye (Zombie NIR^TM^ Fixable Viability Kit), followed by a surface anti-CD56 antibody staining, and fixed with 100 µl/well Fixation Solution (BD Biosciences). Next cells were washed twice with 200 µl Permeabilization Buffer (BD Biosciences) and centrifuged (5 min, 500g). Subsequently cells were stained with IFN-γ-APC and TNF-α-PE (BD Biosciences) for 30 min at room temperature. In the last step, cells were centrifuged (5 min, 500g), resuspended in 200 µl PBS and analyzed using BD FACSCanto II (BD Biosciences).

### NK natural cytotoxicity and ADCC

K562 cells were stained with CellTrace™ CFSE Cell Proliferation Kit (Thermo Fisher Scientific) for 10 min at 37°C. K562 cells were seeded on a U-bottom 96-well at the density of 0.5 × 10^5^ K562 per well and mixed either with 0.25 × 10^6^ NK cells (E:T ratio 5:1) or 0.1 × 10^6^ NK cells (E:T ratio 2:1). For the experiments with conditioned medium, cells were seeded in the 100% conditioned medium. In some experiments, NK cells were incubated with ammonia 4, 24 or 48 hours before assay as indicated in the figure. For ADCC experiments RTX at a final concentration of 100 µg/ml or daratumumab at a final concentration of 1 µg/ml was added. The plate was incubated for 4h at 37°C. Subsequently, cells were stained with 4 µg/ml propidium iodide (PI) and viability was analyzed with BD FACSCanto II (BD Biosciences) as a percentage of propidium iodide-positive CFSE-positive cells.

### Real-time cell analysis (RTCA) cytotoxic assay

The xCELLigence RTCA system (ACEA Biosciences) was used to monitor the viability of the MCF-7 cells during the coincubation with the NK cells and NH_4_Cl. MCF-7 cells were seeded on 16-well E-Plates (ACEA Biosciences) at a cell density 3 × 10^4^ per well in 150 μl of the DMEM medium and monitored for 24h. Next, the medium from the E-Plates with target cells was removed and replaced with NK cells added at E:T ratio 2:1 together with trastuzumab (final concentration 10 µg/ml) and ammonium chloride. The cells were monitored with the RTCA system for the next 12-20h. The analysis of the results was performed using RTCA Software Pro (ACEA Biosciences) and are presented as a normalized cell index.

### Luciferase-based killing assay

Luciferase (luc)-based assays were used to determine cytotoxicity of CD19-CAR-NK92 cells against Raji-luc cells. Target cells were seeded onto U-bottom 96-well black plates at the density of 0.25 × 10^5^ cells per well in 50 µl of DMEM medium. NK cells were added at density 0.25 × 10^5^ (E:T ratio 1:1) in 50 µl of DMEM medium together with 100 µl NH_4_Cl (for the final concentration in range of 1.25 mM-10 mM). Plate was incubated for 18h at 37°C in a humidified atmosphere with 5% CO_2_. The following day 100 μl of each sample was transferred to a white 96-well plate and 100 μl of the the mix of Bright-Glo^TM^ Luciferase Assay (Promega) was added to each well. The plate was incubated for 5 minutes in darkness, the bioluminescence signal was detected using Victor Plate Reader (PerkinElmer).

### Conjugates assay

A total of 6 × 10^6^ NK cells were stained with CellTracker™ Deep Red Dye (Thermo Fisher Scientific) for 10 min at 37°C. At the same time 12 × 10^6^ K562 cells was stained with CellTrace™ CFSE Cell Proliferation Kit (Thermo Fisher Scientific) for 10 min at 37°C. Next cells were washed thoroughly in PBS and resuspended in 3 ml of cell medium. NK cells and K562 were divided and incubated with NH_4_Cl (Sigma Aldrich) for 2h at 37°C. This was followed by co-incubation of NK cells (0.375 × 10^6^) with K562 cells (0.75 × 10^6^) in a total of 100 µl. The reaction was stopped by brief vortexing and addition of 100 µl 4% paraformaldehyde at different time points (0, 5, 15, 30 and 45 min). Cells were analyzed immediately using a FACSCalibur (BD Biosciences) and conjugates were determined as double-positive events.

### Detachment assay

NK and K562 cells were stained using Deep Red and CFSE dyes respectively, as described before. NK cells and K562 were divided and incubated with NH_4_Cl (Sigma Aldrich) for 2h at 37°C. 0.6 × 10^6^ NK cells were mixed with 1.2 × 10^6^ K562 cells in 600 µl total volume in 50 ml falcons, centrifuged (300g, 1 min) and coincubated for 30 min at 37°C to form initial conjugates. Then 2.5 ml of cell medium was added to each falcon and cells were incubated while rotating at 37°C, allowing the NK cells to detach but preventing the formation of new conjugates. At different time points (0, 10, 20, 40, 60 and 90 min) reaction was stopped by addition of 2.5 ml of 4% paraformaldehyde. Cells were analyzed immediately using a FACSCalibur (BD Biosciences) and conjugates were determined as double-positive events.

### Degranulation assay

NK cells (0.2 × 10^6^) and K562 cells (0.1 × 10^6^, E:T ratio 2:1) per well of U-bottom plate were coincubated for 4h at 37°C in the presence of NH_4_Cl and anti-CD107a antibody. Subsequently, cells were stained with viability dye (Zombie NIR^TM^ Fixable Viability Kit) and NK cells’ surface markers antibodies (anti-CD56 and anti-CD3). After that, cells were analyzed using BD FACSCanto II (BD Biosciences).

### Serial degranulation assay

K562 cells were stained with CellTrace™ CFSE Cell Proliferation Kit (Thermo Fisher Scientific) for 10 min at 37°C. Next cells were seeded 0.2 × 10^6^ NK and 0.1 × 10^6^ K562 (E:T ratio 2:1) per well and coincubated for 2h at 37°C in the presence of NH_4_Cl and anti-CD107a-PE antibody. After that time cells were washed and fresh NH_4_Cl was added together with anti-CD107a-BV421 antibody and incubated for 4h at 37°C. Cells were analysed using BD FACSCanto II (BD Biosciences). Serial degranulation was defined as double-positive events (CD107a-PE^+^ and CD107a-BV421^+^).

### Lentiviral modification of NK-92 cells with CARs

NK-92 cells were transduced with pSEW-CD19 (FMC63)-CH3-IgG1-CD28-CD3z (CD19 CAR) or atezolizumab-based anti-PD-L1 CAR ^68,69^. To produce CD19 or PD-L1 CAR viral particles, HEK-293T cells were seeded at 10 cm plates and transfected using a polyethyleneimine (PEI) transfection protocol simultaneously with CAR-encoding plasmids, VSV-G envelope expressing plasmid pMD2.G (RRID: Addgene_12259) and lentiviral packaging plasmid psPAX2 (RRID: Addgene_12260). After 48 h, the lentivirus-containing supernatant was harvested, filtered through a 0.45 μm pore size filter, and concentrated by overnight centrifugation at 2 500 × g at 4°C. The culture medium from the NK-92 cells was replaced with concentrated lentiviral supernatant supplemented with 15 µg/ml protamine sulfate and 6 µM BX-795. After 1 h of spinoculation (750 × g at 25°C), the NK-92 cells were kept overnight in an incubator. The next day, the viral supernatant was replaced with the fresh portion of complete culture X-VIVO™ 20 medium (Lonza) supplemented with 200 U/ml IL-2 (PeproTech). The CAR expression on the surface of the NK-92 cells was evaluated by flow cytometry 48-72 h after transduction.

### ELISA

NK cells together with K562 cells were treated with NH_4_Cl for 4h at 37°C. Subsequently cells were extensively mixed, centrifuged, and supernatant were collected. It was analysed using Human Perforin ELISA kit (MABTECH) according to the manufacturer’s protocol and measured using PerkinElmer Multimode Plate Reader EnVision. This kit uses two types of anti-perforin antibodies, Pf-344 that recognize an epitope after perforin monomers undergo pore formation, and Pf-80 that recognize a conformational epitope accessible at acidic but not neutral pH.

### Western blotting

The total of 1 × 10^6^ NK cells per well were seeded on a 6-well plate and incubated for 4h at 37°C in the presence of NH_4_Cl. Afterward, cells were suspended in ice cold lysis buffer (150 mM NaCl, 1% Triton X-100, 50 mM Tris-HCl pH 8.0) supplemented with protease inhibitors (Roche) and incubated on ice for 30 min. The lysates were centrifuged for 30 min at 12,000g at 4°C, supernatants were transferred to new tubes. Total protein concentration was measured using a bicinchoninic acid (BCA) assay. 5 to 20 µg of the protein lysate was mixed with 5-times concentrated loading Laemmli buffer (0.125 M Tris pH 6.9, 4% SDS, 10% DTT, 20% glycerol) and samples were boiled for 5 min at 95°C. Proteins were separated using 4-12%Bis-Tris NuPAGE gels (Life Technologies). They were next transferred to a PVDF membrane which was blocked with 5% nonfat dry milk for 1h. Membranes were incubated overnight at 4°C with perforin antibodies. Total perforin was detected with Abcam (#ab47225) antibody, mature and immature forms of perforin were detected with Mabtech (#3465-6-250) antibody. Subsequently, membranes were washed and incubated with HRP-conjugated secondary antibodies for 2h at room temperatures. ß-actin antibody (#A222, Sigma Aldrich) was used as a loading control. For protein detection SuperSignal West Pico PLUS substrate (#34580, Thermo Fisher Scientific) was used. The signal was detected with ChemiDoc Imaging System (Bio-Rad Laboratories).

### Live-cell imaging with reporter cells

To investigate killing mechanisms 0.2 × 10^5^/ml HeLa-CD48 cells that stably express NES-ELQTD-GFP-T2A-NES-VGPD-mCherry and CD48 were seeded onto a silicon-glass microchip and left to adhere overnight. Fluorescent reporters allow to measure granzyme B (NES-RIEADS-mCherry) and caspase-8 (NES-ELQTD-GFP) activity shortly after NK cells attached to the target cell. The following day 0.1 × 10^4^/ml NK cells were added to the microchip together with NH_4_Cl and placed into the incubation chamber (37°C, 5% CO_2_). Time-lapse live-cell microscopy was immediately started using a Zeiss Axio Observer Z1 7 microscope. EGFP and mCherry were excited using the Colibri 7 LED-module 475 (filter set 90 HE LED) and 567 (filter set 91 HE LED), respectively. Images were acquired every 3 min for 15–17h using a Hamamatsu ORCA-flash 4.0 camera. Images were analyzed using ImageJ software.

### Serial killing

To evaluate NK cells’ serial killing ability, 0.5 × 10^6^/ml K562 cells were seeded onto a silicon-glass microchip, together with viability dye 2.5 μM SYTOX^TM^ Blue (Thermo Fisher Scientific). Subsequently, 0.375 × 10^6^/ml NK cells were added together with NH_4_Cl and placed into the incubation chamber (37°C, 5% CO_2_). The analysis required tracking a randomly selected single NK cell and assessing its interaction with a target cell. Target cells’ viability was determined based on the intensity of fluorescence dye SYTOX^TM^ Blue, which easily penetrates cells with damaged cell membrane and stain cell nuclei. Time-lapse live-cell microscopy was immediately started using a Zeiss Axio Observer Z1 7 microscope. Images were acquired every 3 min for 15–17h using a Hamamatsu ORCA-flash 4.0 camera. Images were analyzed using ImageJ software. Results were presented as a diagram showing the serial killing of NK cells.

### Microscopy

For the analysis of the lysosomal content, NK cells were incubated with ammonia (ammonium chloride) for 12 hours. Then, LysoTracker Deep Red (Thermo Fisher Scientific) was added for 30 minutes at final concentration of 50 nM. Nuclei were stained with Hoechst (H1399; Thermo Fisher Scientific) with the final concentration of 5 μg/mL. Cells were imaged using Opera Phenix live cell microscopy (PerkinElmer). Images were analyzed using Harmony 4.9 Software (PerkinElmer). At least 10 fields were analyzed for each of 4 NK cells donors. For LAMP1 and CD63 analysis, NK cells were incubated with ammonia (ammonium chloride) for 12 hours. Then, cells were rinsed twice for 5 min with ice-cold PBS and fixed with 3% paraformaldehyde in PBS for 12 min at room temperature, followed by a simultaneous permeabilization with 0.1% (w/v) saponin and blocking with 0.2% (w/v) fish gelatin in PBS for 10 min. They were further incubated with appropriate primary and secondary antibodies in 0.01% (w/v) saponin and 0.2% fish gelatin in PBS for 30 min each. The mouse anti-CD63 antibody (mAb, #H5C6) developed by J.T. August/J.E.K. Hildreth was obtained from the Developmental Studies Hybridoma Bank developed under the auspices of the NICHD and maintained by The University of Iowa, Department of Biology, Iowa City, Iowa, USA. Rabbit anti-LAMP1 (L-1418, 1:400) was from Sigma-Aldrich. Secondary antibodies used for immunofluorescence were: Alexa Fluor 488-, 555-, 647-conjugated anti-mouse-IgG and anti-rabbit-IgG were from ThermoFisher Scientific. All secondary antibodies were diluted 1:10,000. Airyscan imaging was performed with a confocal laser scanning microscope ZEISS LSM 800 equipped with Plan-Apochromat 63×/1.40 NA oil objective and an Airyscan detection unit, using Immersol 518 F immersion medium (Zeiss). Detector gain and pixel dwell times were adjusted for each dataset, keeping them at their lowest values to avoid saturation and bleaching effects. ZEN Blue 2.3 (Version 2.3.69.1003) software (Zeiss) was used for image acquisition. The Airyscan processing module with default settings was used to process obtained data. Pictures were assembled in Photoshop (Adobe) with only linear adjustments of contrast and brightness.

### Statistical analysis

Data are shown as means ± SD or means ± SEM, as indicated in the figure legends. GraphPad Prism 9.5.1 (GraphPad Software) was used for statistical analyses. Data distribution was tested using the Shapiro–Wilk test, D’Agostino & Pearson test, and Kolmogorov–Smirnov test. Statistical analyses of three or more groups were compared using one-way or two-way analysis of variance (ANOVA) or Brown–Forsythe ANOVA followed by Tukey’s, Dunnett’s, or Bonferroni’s multiple comparisons test or Kruskal–Wallis’s test followed by Dunn’s multiple comparisons test. Repeated measures ANOVA with Sidak’s or Holm–Sidak’s post hoc tests were used to analyze the differences in paired samples. Statistical analyses of two groups were compared using unpaired t-test, paired t-test, or Mann–Whitney test. Methods of statistical analyses are defined in every figure legend. A *P* value of less than 0.05 was considered as statistically significant. Each experiment was performed in technical duplicates or triplicates. The number of biological replicates for each experiment is mentioned in the figure legends.

## ACKNOWLEDGEMENTS

The research leading to these results has received funding from the European Research Council 805038/STIMUNO/ERC-2018-STG (M.W.). This project was also supported by Norway Grants 2014-2021 through the National Centre for Research and Development (POLNOR/ALTERCAR/0056/2019-00) (M.W.), National Science Centre UMO-2019/35/D/NZ6/03073 (A.G-J). T.M.G. is supported by the Foundation for Polish Science (FNP), A.G-J is supported by the Polish Ministry of Education and Science. We would like to thank Ewa Pieta for excellent technical assistance.

## AUTHORS CONTRIBIUTIONS

J.D. conducted most *in vitro* experiments and performed initial analysis of the data. T.M.G. contributed to some *in vitro* flow cytometry experiments, analyzed the data, prepared figures, and wrote the manuscript. I.B. designed, coordinated and conducted *in vivo* experiments. A.K. performed *in vitro* experiments on the influence of ammonia on ADCC. K.F. designed and conducted *in vivo* experiments. K.M. performed *in vitro* experiments on the influence of ammonia on ADCC and CAR-NK activity. Z.P. optimized and conducted some experiments with TIF and SCF isolation. M.G. performed *in vitro* experiments with the conditioned medium. A.G-J. performed preliminary *in vitro* experiments on the role of ammonia in the regulation of NK cells activity. L.K.P. performed and analyzed experiments on the mechanisms of killing by NK cells and analysis of serial killing. K.J. performed microscopy imaging of NK cells and analyzed lysosomal content in NK cells. M.J. performed analysis of ammonia accumulation in NK cells. D.U. supervised experiments on the mechanisms of killing by NK cells. M.B. edited the manuscript. C.W. supervised the study on NK cell serial killing. M.M supervised the study on the visualization of the lysosomes. M.W. conceived, designed, and supervised the study, provided funding and resources, and supervised and edited the manuscript. All authors provided critical feedback, reviewed and approved the final manuscript.

