## Supplemental Figures for "Ammonia inhibits antitumor activity of NK cells by decreasing mature perforin"

### SUPPLEMENTARY FIGURES

### Ammonia inhibits antitumor activity of NK cells by impairing perforin maturation

Joanna Domagala et al.

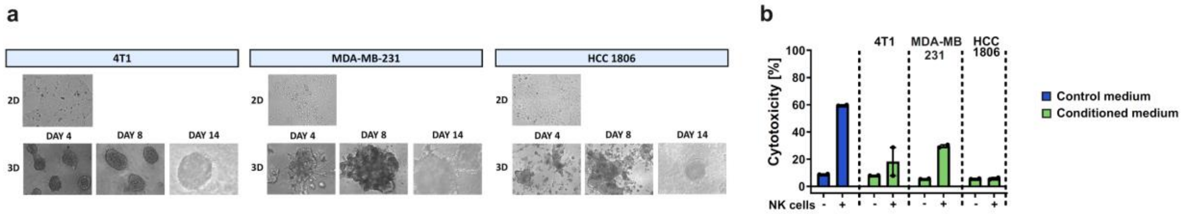


**Supplementary Fig. 1. 3D cancer culture-conditioned medium suppresses cytotoxicity of NK cells**

**a**, Microscopy photographs of 4T1, MDA-MB-231 and HCC 1806 cells cultured in conventional conditions (2D) and three-dimensional culture (3D). Conditioned medium was collected on the 14$^{th}$ day after 48 hours of culture. **b**, Natural cytotoxicity of NK cells against K562 cells in the presence of control medium and breast cancer 3D culture-conditioned medium. Data from a representative experiment. K562 cells were stained with CFSE and incubated with NK cells in a medium conditioned by indicated cells. Cytotoxicity was assessed after 4 hours using flow cytometry and presented as percentage of propidium iodide-positive CFSE-positive (K562) cells.

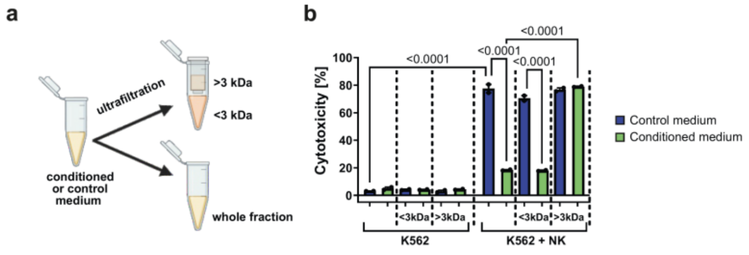


**Supplementary Fig. 2. Low molecular fraction of conditioned medium suppresses cytotoxicity of NK cells**

**a**, Raji-cells conditioned medium was divided using an ultrafiltration method into a low molecular fraction (< 3kDa) and a higher molecular fraction (> 3kDa). **b**, Natural cytotoxicity of NK cells against K562 cells in the presence of the whole fraction, low molecular fraction (<3 kDa) and higher molecular fraction (> 3 kDa) of control and Raji cells-conditioned medium. Data from a representative experiment. K562 cells were stained with CFSE and incubated with NK cells in a medium conditioned by indicated cells. Cytotoxicity was assessed after 4 hours using flow cytometry and plotted as percentage of propidium iodide-positive CFSE-positive (K562) cells.

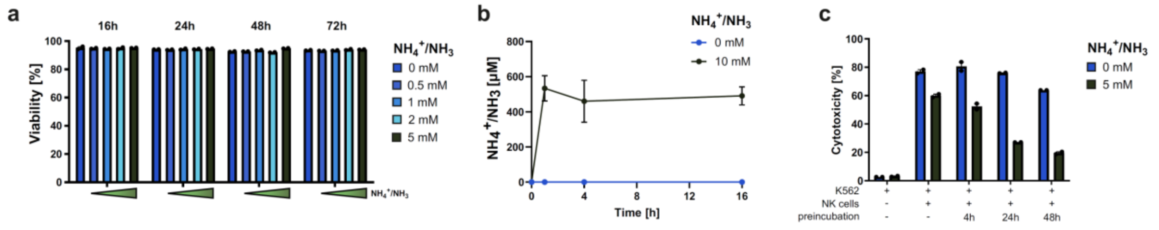


**Supplementary Fig. 3. Ammonia accumulates in NK cells, affects their cytotoxic potential but not their viability.**

**a**, Viability of NK cells incubated with different concentrations of ammonia (ammonium chloride) for 16, 24, 48 and 72 hours was assessed using propidium iodide staining and flow cytometry (n=2). **b,** The concentration of ammonia in NK cells after incubation with ammonia (ammonium chloride) for 30 min, 4 and 16 hours (n=3). After the indicated time, cells were washed two times with PBS and lysed. **c**, Natural cytotoxicity of NK cells against K562 cells in the presence of ammonia (ammonium chloride) (n=2). Cytotoxicity was assessed after 4 hours using flow cytometry and determined as percentage of propidium iodide-positive CFSE-positive (K562) cells. NK cells were preincubated with ammonia for 4, 24 or 48 hours, as indicated in the figure.

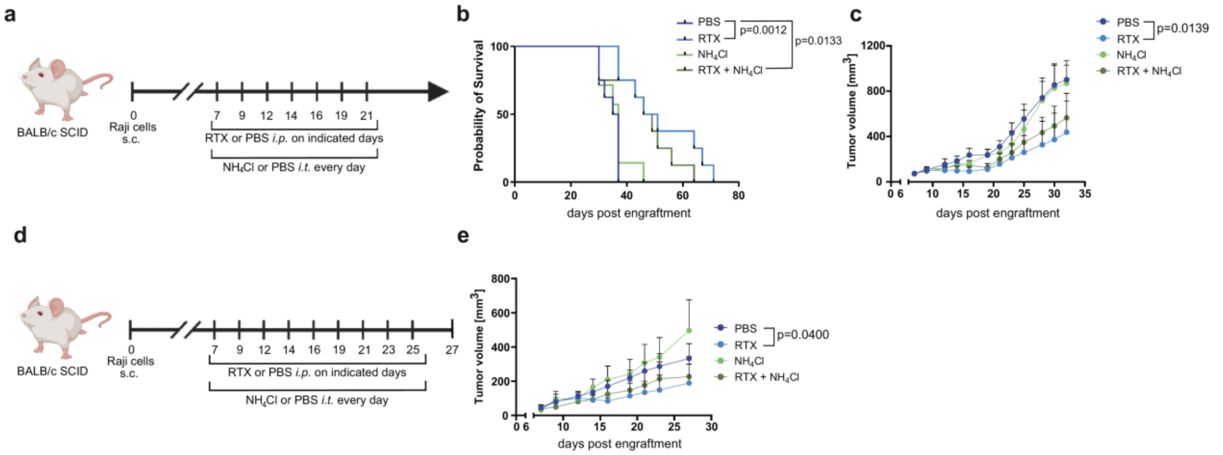


**Supplementary Fig. 4. Ammonium chloride's impact on the efficiency of rituximab therapy *in vivo***

**a**, In vivo experimental scheme. Raji cells were injected into BALB/c mice. 7 days after injection, mice started receiving rituximab (RTX) or PBS intraperitoneally (*i.p.*) and were injected intratumorally (*i.t.*) with either ammonia (ammonium chloride) or PBS for 14 days. **b-c**, Survival (**b**), and tumor growth (**c**) of mice in each group (n=5 mice/group). **d**, In vivo experimental scheme. Raji cells were injected into BALB/c mice. 7 days after injection, mice started receiving RTX or PBS (*i.p.*) and were injected *i.t.* with either ammonia (ammonium chloride) or PBS for 18 days. **e,** the graph shows the tumor growth (**c**) of mice in each experimental group (n=5 mice/group).

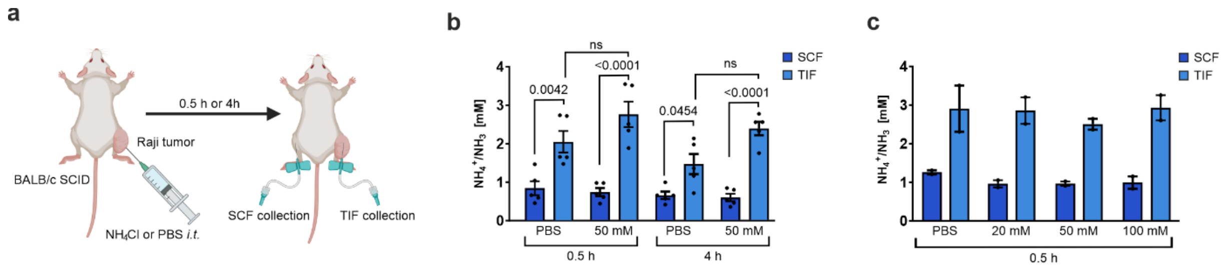


**Supplementary Fig. 5. Ammonia is rapidly excreted or metabolized in TME**

**a**, *In vivo* experimental scheme. Raji cells were injected into BALB/c mice. 7 days after injection, mice were injected *i.t.* with either ammonia (ammonium chloride) or PBS. After 30 min or 4 hours TIF and SCF were collected followed by an ammonia measurement. **b-c**, Concentration of ammonia in TIF and SCF isolated from Raji tumors after *i.t.* administration of ammonia (ammonium chloride) or PBS (**b,** n=5; **c**, n=2).

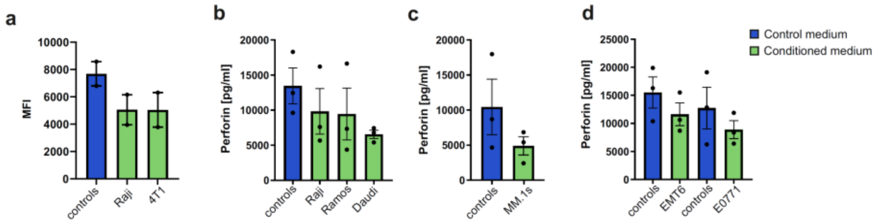


**Supplementary Fig. 6. Conditioned medium decreases expression and secretion of perforin by NK cells**

**a**, The level of perforin detected in NK cells incubated with Raji- or 4T1-conditioned medium determined by intracellular staining using anti-perforin antibody (δG9 clone) and flow cytometry (n=2). **b-d,** The concentration of extracellular perforin secreted by NK cells in response to contact with target cells (K562) in **b**, lymphoma cells-conditioned medium (n=3), **c,** multiple myeloma cells-conditioned medium (n=3), and **d,** breast cancer cells-conditioned medium (n=3).

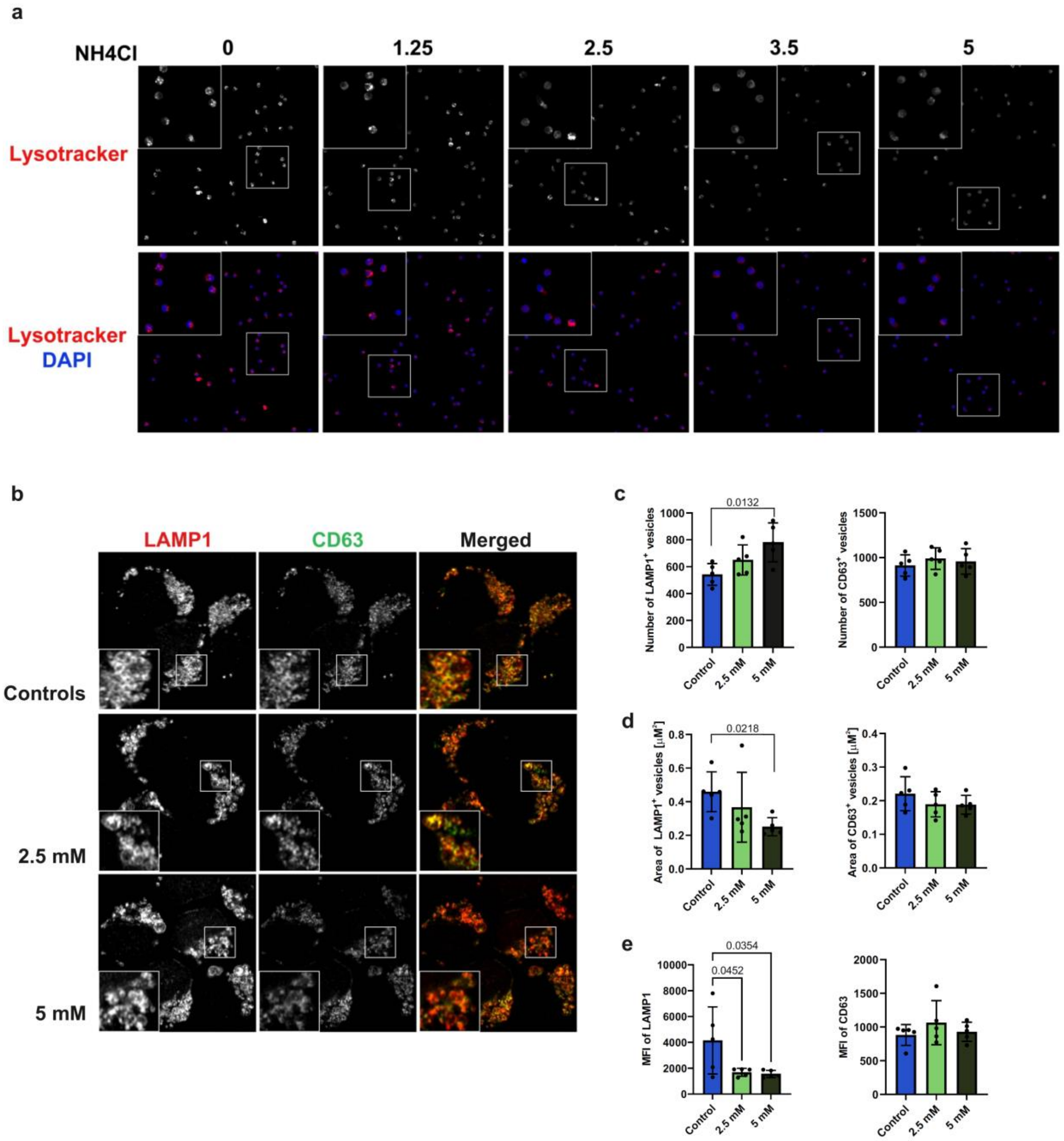


**Supplementary Fig. 7. Ammonia increases the number of LAMP1$^+$ vesicles but decreases their volume and LAMP1 level**

**a**, The lysosomal content of NK cells incubated with ammonia (ammonium chloride) for 12 hours. Acidic organelles were stained using a LysoTracker and live imaged using Opera Phenix cell microscopy (n=4). *P* values were calculated using two-way ANOVA with Tukey's post hoc test. Data show individual values and means ± SEM. *n* values are the numbers of biological replicates in *in vitro* experiments. **b,** LAMP1 and CD63 staining of NK cells incubated in different concentrations of ammonia (ammonium chloride). Cells were incubated with ammonia for 4 hours, followed by antibody staining and imaging using ZEISS LSM 800 with Motiontracking. **c-d,** Number (**c**) and area (**d**) of LAMP1$^+$ and CD63$^+$ vesicles in a single NK cell incubated with different concentrations of ammonia (ammonium chloride) (n=5). **e,** Mean fluorescence intensity (MFI) of LAMP1 and CD63 staining in a single NK cell incubated with different concentrations of ammonia (ammonium chloride) (n=5).
